# The force loading rate drives cell mechanosensing through both reinforcement and fluidization

**DOI:** 10.1101/2021.03.08.434428

**Authors:** Ion Andreu, Bryan Falcones, Sebastian Hurst, Nimesh Chahare, Xarxa Quiroga, Anabel-Lise Le Roux, Zanetta Kechagia, Amy E.M. Beedle, Alberto Elósegui-Artola, Xavier Trepat, Ramon Farré, Timo Betz, Isaac Almendros, Pere Roca-Cusachs

**Affiliations:** Institute for Bioengineering of Catalonia (IBEC), the Barcelona Institute of Technology (BIST), 08028 Barcelona, Spain; Universitat de Barcelona, 08036 Barcelona, Spain; Universitat Politècnica de Catalunya (UPC), Campus Nord, Carrer de Jordi Girona, 1, 3, 08034 Barcelona; Institute of Cell Biology, Center of Molecular Biology of Inflammation (ZMBE), University of Münster, Von-Esmarch-Straße 56, 48149 Münster; Department of Physics, King’s College London, Strand, WC2R 2LS London, United Kingdom; Harvard John A. Paulson School of Engineering and Applied Sciences, Harvard University, 58 Oxford St., Cambridge, MA, 02138 USA; Wyss Institute for Biologically Inspired Engineering, 3 Blackfan Circle, Boston, MA, 02115 USA; Institució Catalana de Recerca i Estudis Avançats (ICREA), Passeig de Lluís Companys, 23, 08010 Barcelona; CIBER de Enfermedades Respiratorias, Madrid; Institut d’Investigacions Biomèdiques August Pi Sunyer, Barcelona

## Abstract

Cell response to force regulates essential processes in health and disease. However, the fundamental mechanical variables that cells sense and respond to remain unclear. Here we show that the rate of force application (loading rate) drives mechanosensing, as predicted by a molecular clutch model. By applying dynamic force regimes to cells through substrate stretching, optical tweezers, and atomic force microscopy, we find that increasing loading rates trigger talin-dependent mechanosensing, leading to adhesion growth and reinforcement, and YAP nuclear localization. However, above a given threshold the actin cytoskeleton undergoes fluidization and softens, decreasing loading rates and preventing reinforcement. By stretching rat lungs *in vivo*, we show that a similar phenomenon occurs at the organ level. Our results show that cell sensing of external forces and of passive mechanical parameters (like tissue stiffness) can be understood through the same mechanisms, driven by the properties under force of the mechanosensing molecules involved.

## Introduction

Cells are constantly subjected to forces transmitted through tissues^1, 2^, which regulate major processes in health and disease^3, 4^. Despite this importance, the fundamental mechanical variables that cells sense and respond to are not fully understood, and have been a matter of intense debate^5–7^. Mechanosensing molecules such as the integrin-actin adaptor protein talin respond to specific values of applied force^8^, but it has been suggested that cells respond not directly to force but to the associated deformations exerted on the extracellular matrix (ECM)^5, 9^ or even to a combination of force and deformation^6, 10^. However, in most scenarios forces and deformations are highly dynamic^11–14^, also because physiological ECMs are typically viscoelastic (and therefore have time-dependent responses)^15, 16^.Thus, a sensing system based only on given fixed magnitudes of force or deformation may not be effective. Alternatively, cells could be sensitive to force dynamics per se. Specifically, force sensitive molecular events such as bond rupture^17, 18^ or protein unfolding^19^ have long been predicted and measured to depend on the rate of force application, known as the loading rate. This dependency is in fact an implicit underlying assumption of the molecular clutch theory, which has been employed to model how cells generate and transmit forces to sense passive mechanical factors such as ECM rigidity^20, 21^, viscosity^22^, or ligand density^23^. In this framework, cells generate forces through actomyosin contraction, which results in a retrograde flow of actin from the cell edge to the cell centre. As actin is connected to the ECM via integrins, this retrograde flow pulls on and deforms the substrate. ECM mechanics strongly influence the resulting loading rate, which is the product of the deformation speed times the effective stiffness of the substrate. Experimentally, this effect can be seen for instance by observing the dynamics of cell pulling of micropillars of different stiffness^10^. Changes in loading rates then enable ECM mechanosensing by affecting molecular events such as integrin-ECM binding or talin unfolding^8, 20^. This hypothesis is attractive in that it relates cell sensing of passive ECM mechanical factors to the sensing of directly applied forces, in a unified mechanism. It is also consistent with the well-known frequency dependence of cell mechanoresponse in many different systems^24–27^. However, whether the loading rate (rather than other static or dynamic mechanical variables) is a driving parameter of mechanosensing, is unknown.

## Results

### The cell stretch rate drives mechanosensing in a biphasic manner

To start exploring the role of loading rate, we seeded mouse embryonic fibroblasts on very soft (0.6 kPa in rigidity) fibronectin-coated polyacrylamide gels. In these conditions, we measured actin retrograde flows via time-lapse imaging of lifeact-transfected cells, and mechanosensing via immunostaining of two well-known mechanosensitive features: paxillin-containing cell-matrix adhesions, and the nuclear translocation of the transcriptional regulator YAP^28, 29^. Retrograde flows were of ∼50 nm/s (Fig. 1a,b and Supplementary Video 1) and mechanosensing was not triggered, as cells showed largely cytosolic YAP and only small punctate paxillin-containing adhesions (Fig. 1c). This is consistent with previously reported phenotypes of cells on soft substrates, and is indicative of a fast flow caused by low adhesion (and thereby low actin attachment) to the ECM^20^. Consistently, actin flows of cells on stiff substrates but measured at the very edge of lamellipodia (where adhesion is still very low) were of a similar magnitude (Fig. 1a,b and Supplementary Video 2).

**Fig. 1:**
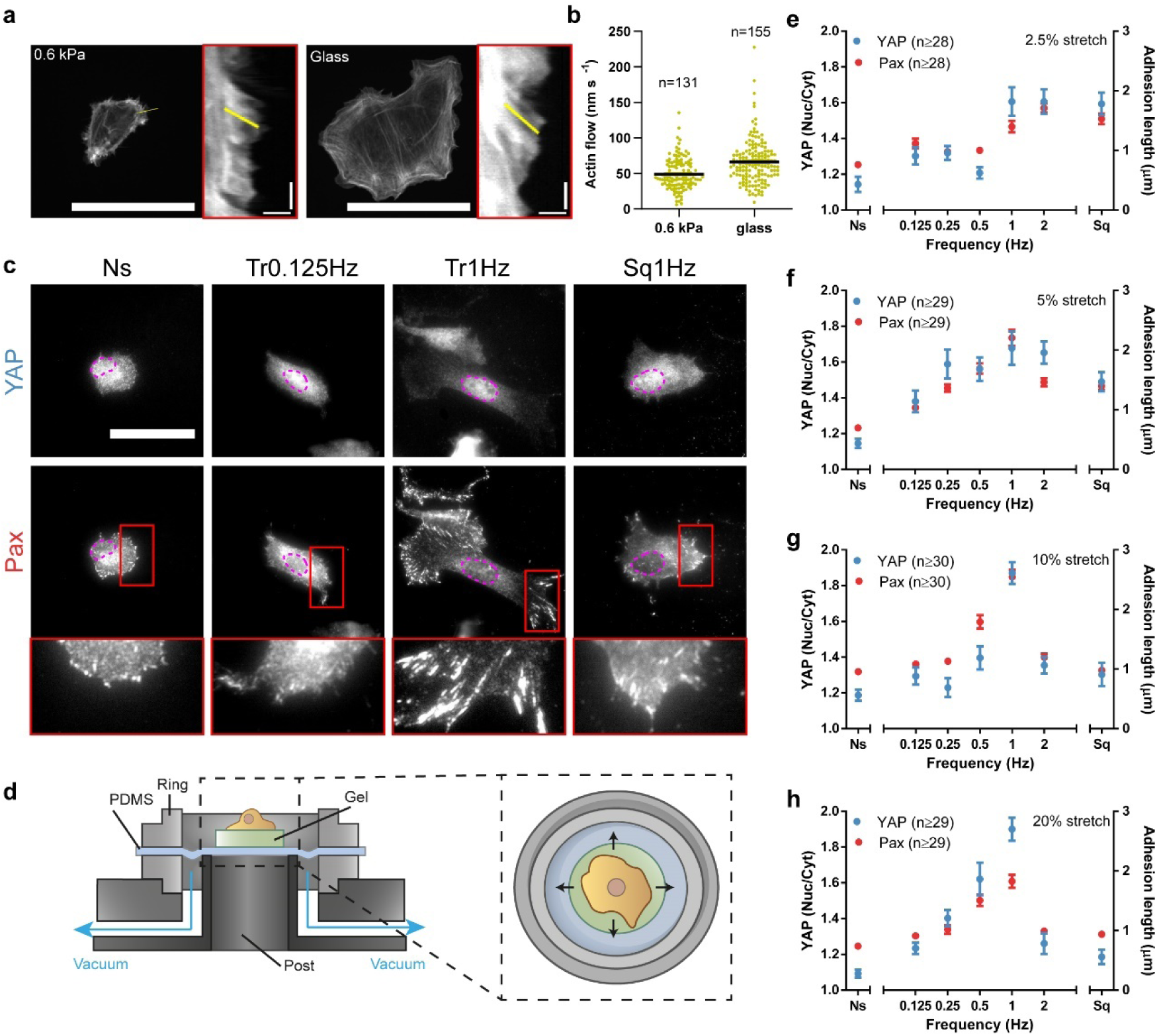
The rate of cell stretch drives mechanosensing in a biphasic manner. a. Cells transfected with LifeAct-GFP and plated on polyacrylamide gels of 0.6 kPa or on glass. Insets are kymographs showing the movement of actin features along the lines marked in yellow. **b** Actin retrograde flow of cells cultured on 0.6 kPa gels or glass. n numbers are traces. **c**, YAP and paxillin stainings of cells stretched by 10% using the setup using triangular (Tr) and square (Sq) signals at different frequencies. Ns, non-stretched cells. In YAP images, magenta outlines indicate the nucleus. In paxillin images, areas circled in red are shown magnified at the right. **d**, Illustration of the stretch setup. **e-h**, Quantifications of YAP nuclear to cytoplasmic ratios and paxillin focal adhesion lengths for cells stretched at 2.5 % (e), 5% (f) 10% (g), and 20% (h). Results are shown for non-stretched cells (Ns), cells stretched with triangular signals at different frequencies, and cells stretched with a square signal at 1 Hz. The effects of frequency were significant for both YAP and paxillin in all panels (p<0.0001). The effect of square versus triangular 1 Hz signals was significant for paxillin at 5% stretch (p=0.0025) and for both YAP and paxillin for 10% and 20% stretch (p<0.0001). n numbers are cells. Scale bars are 50 μm in cells and 2 μm/40 sec in kymographs (x/y axes). Data are shown as mean ± s.e.m.

We then used two ways to increase the loading rate, considering that it is the product of deformation speed times substrate stiffness. First, we increased the loading rate indirectly, by increasing substrate stiffness. In this case, mechanosensing (as indicated by changes in YAP nuclear localization) was first observed when rigidity was increased over 5-fold to 3.4 kPa (Supplementary Fig. 1a). Second, we increased the loading rate directly, by externally applying stretch (a stimulus well known to trigger mechanosensing^24–27)^ to cells seeded on soft 0.6kPa gels using a previously described device (Fig. 1d). From a given amplitude and frequency of applied stretch, we can estimate the corresponding cell deformation speeds. To this end, we consider the average spreading diameter of cells (∼20 μm), and assume that cell-substrate attachment and force transmission occurs largely at the cell periphery, where focal adhesions were mostly located. Using this approach, applying a very mild stretch (2.5% biaxial stretch, applied cyclically with a triangular 0.125 Hz wave for 1 h) leads to a deformation speed of ∼60 nm/s, of the same order of magnitude than internally generated actomyosin flows in low adhesion conditions. Consistently, this signal had only a very small effect on YAP or paxillin mechanosensing responses (Fig. 1e). However, when we increased deformation speeds by changing stretch frequency (and not amplitude), a clear response was observed above 1 Hz both for YAP (by progressively localizing in the nucleus) and for adhesions (by growing from punctate to the larger, elongated structures known as focal adhesions, Fig. 1e). Compared to the signal at 0.125 Hz, this represents an increase of 4-8 fold, again consistent with stiffness results. Whereas this equivalence is only approximate, and the magnitude of response was different in both cases (higher with stiffness than with stretch), the match in order of magnitude support a role of the loading rate. Importantly, as previously reported for stiffness^20^, the mechanosensing response to stretch was mediated by the mechanosensitive protein talin^20^, since its knock-down eliminated both YAP and paxillin responses (Supplementary Fig. 1d-g).

We then repeated the frequency sweep by applying different amounts of stretch, from 2.5% to 20% (Fig. 1e-h, and Supplementary Fig. 2a-h). As expected from a role of loading rate, progressively increasing the stretch amplitude led to higher responses at most frequencies. However, for stretch amplitudes above 5%, increasing the frequency only increased response up to a point: above 1 Hz, fast stretching failed to trigger adhesion growth or YAP nuclear translocation (Fig. 1c,e-h). This was not due to cells detaching from the substrate, since spreading areas did not decrease (Supplementary Fig. 1b). To further verify the role of the stretch rate and to decouple it from that of stretch frequency, we stretched cells at a frequency of 1 Hz, but instead of applying a progressive triangular signal, we applied stretch as fast as possible. This led to a quasi-square signal where the stretch rate more than doubled with respect to the corresponding triangular signal, and was thereby somewhat above that of the triangular 2 Hz signal (Supplementary Fig. 2b,h). Accordingly and for all stretch amplitudes, applying a square rather than triangular 1 Hz signal led to similar results than those obtained at the 2 Hz triangular signal (Fig. 1c,e-h). Importantly, this observed biphasic response was also observed in cell proliferation rates (Supplementary Fig. 1c), a well-known downstream effect of mechanosensing and YAP^28^. The biphasic response was also generalizable to other cell types (lung endothelial and epithelial cells, supplementary fig. 3).

To understand the decreased response at high frequencies, we hypothesized that it could be caused by the well-described phenomenon known as fluidization, by which both cells and actin gels soften when submitted to high stretch amplitudes^30–33^ or rates^34^. Fluidization is caused by a partial disruption of the cytoskeleton, that is explained not by a single molecular event but by broad effects on the entire cytoskeletal network^30^. Of note, in accordance to existing literature on stretch responses^32, 33^ here we refer to fluidization simply to mean cytoskeletal softening, and not the more precise rheological definition which involves both softening and an increased viscous-like behaviour^30, 35^. To assess this hypothesis, we evaluated cytoskeletal organization in the different conditions.

Cells stretched at very low or high rates (1 Hz square signal) showed a largely unstructured cytoskeleton (Fig. 2a). Accordingly, their actin anisotropy (a quantification of the degree of alignment and organization of actin fibres^36^) was low, at the level of unstretched cells (Fig. 2b). In contrast, cells stretched by 10% with a 1 Hz triangular signal, where focal adhesions were largest, formed clear actin stress fibres connected to adhesions (Fig. 2a), and exhibited high values of actin anisotropy (Fig. 2b). Then, we stretched cells treated with Jasplakinolide, a drug which stabilizes actin fibres by preventing their depolymerisation^37^. Consistent with a role of cytoskeletal fluidization, Jasplakinolide treatment rescued the response to high stretch rates (1 Hz square signal), in terms of stress fibre formation (Fig. 2a), actin anisotropy (Fig. 2b), and focal adhesion formation (Fig. 2c).

**Fig. 2:**
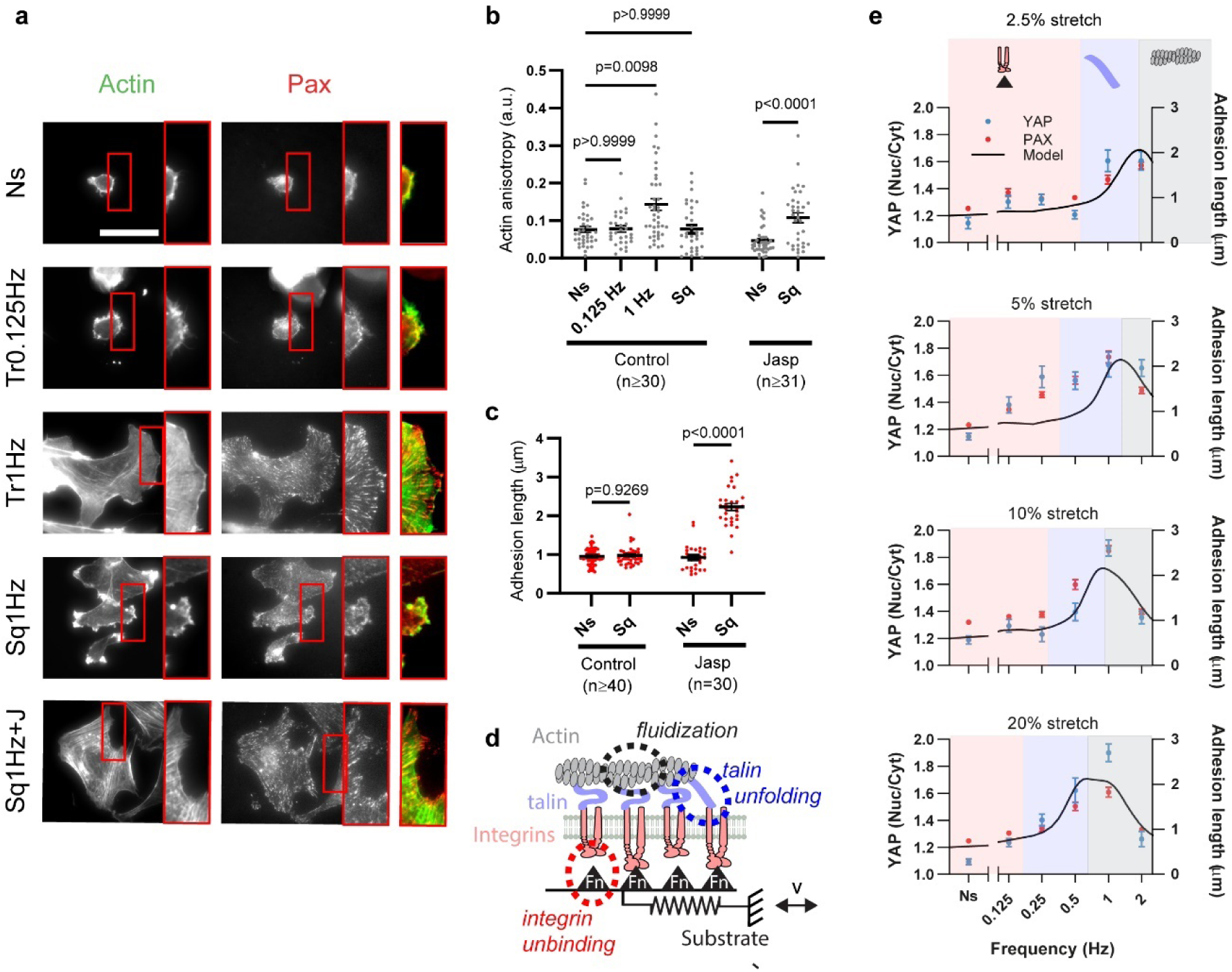
A molecular clutch model considering mechanosensing and fluidization predicts the response to stretch. **a**, Actin and paxillin stainings for cells either not stretched or stretched by 10% with triangular signals (0.125 Hz, 1 Hz) or square signals (1 Hz, with or without Jasplakinolide treatment). Areas circled in red are shown magnified at the right of each image, and shown as a merged image (actin, green, paxillin, red). Scale bar is 50 µm. **b, c** corresponding quantifications of actin anisotropy (b) and adhesion length (c) for control cells, and cells treated with Jasplakinolide. n numbers are cells. **d**, Cartoon of computational clutch model (see methods). The model considers a relative speed of movement v between cell and substrate (given by stretch). The substrate is represented by an elastic spring with binding sites to integrins (via fibronectin, Fn triangles), which in turn connect to talin and actin. As stretch applies forces, these can lead to integrin unbinding, talin unfolding, or cytoskeletal fluidization. **e**, Model predictions (black line) overlaid on experimental YAP and paxillin results from figure 1. The only parameter changing between simulations is stretch amplitude (1.5, 2, 2.5 and 3 µm for 2.5, 5, 10, and 20% stretch). Areas shaded in pink, blue, and gray show the regions dominated respectively by integrin unbinding, talin unfolding, and fluidization. Data are shown as mean ± s.e.m.

### A clutch model considering how the loading rate affects reinforcement and fluidization explains the results

Our results can be explained by a role of loading rate, affecting both reinforcement and fluidization. To assess how these different factors are related, we developed a computational clutch model based on our previous work^20^. This model considers progressive force application to links between actin, talin, integrins, and fibronectin (i.e., “clutches”). Then, it considers how talin unfolding and integrin-fibronectin unbinding depend on force (based on experimental single molecule data^38, 39^). If talin unfolds before the clutch disengages from the substrate through integrin unbinding, we assume that there is a mechanosensing reinforcement event, which leads to integrin recruitment (i.e., adhesion growth in experiments). As a modification from our previous model, here we introduced that i) force on clutches does not arise from actomyosin contractility, but from externally imposed periodic stretch, and ii) the clutch can be disengaged not only by integrin unbinding, but also by actin cytoskeleton disruption (i.e., fluidization) above a threshold force. Importantly, the model does not assume any dependence on loading rate per se. Rather, increasing loading rates simply reduce the amount of time required to reach a given force level, thereby increasing the likelihood that high forces will be reached before clutch disengagement. Thus, different loading rates means that different force regimes are reached, which may differently affect the molecular events considered (integrin unbinding, talin unfolding, or cytoskeletal fluidization).

By modifying only the parameters of applied stretch frequency and amplitude, the model largely reproduced observed experimental trends (Fig. 2e). For low frequencies, low loading rates mean that forces stayed in a regime where integrin unbinding occurred first, preventing mechanosensing. At higher frequencies higher forces were reached, progressively allowing talin unfolding, and increasing mechanosensing. However, at very high frequencies, the very high forces required for fluidization were reached. By increasing stretch amplitude, loading rates are achieved at lower frequencies, shifting the curves to the left, and effectively making the fluidization regime observable only for high amplitudes. Of note, the fluidization event in the model cannot distinguish between different potential events, such as breaking of actin filaments, or severing of actin crosslinks, for instance. However, the model provided a good fit to the data by assuming a force of about 140 pN, in reasonable agreement with reported experimental values for the breaking of actin filaments^40^.

### The loading rate drives the maturation of single adhesions

Our results and modelling are consistent with a role of loading rate, which would trigger mechanosensing or fluidization depending on its magnitude. To further verify this, we tested some assumptions of our hypothesis and model which could not be addressed through the stretch device. First, effects should be caused by the loading rate and not the cell deformation rate, which also increased with stretch. Second, the effects are expected to occur not only at the global cell scale, but also at the local adhesion scale. To address these questions, we used a previously described optical tweezers setup^41^ (Fig. 3a). We seeded cells transfected with GFP-paxillin on glass, trapped fibronectin-coated 1 μm diameter beads, attached them to the cell surface, and applied forces to cells by displacing the optical trap horizontally with triangular signals of the same amplitude, but different frequencies (Fig. 3b,c, and Supplementary Fig. 4a). Initially, this stimulation led to bead displacements of ∼0.2 μm and applied forces of 10-15 pN, which did not show any significant trend with frequency (Supplementary Fig. 4b-d). With time, force application led to the mechanosensing process known as adhesion reinforcement (Fig. 3b). This was characterized by a progressive reduction in bead displacements (Fig. 3c) and speeds (Fig. 3d), a measure of applied deformation rates. Concomitantly, there was an increase in applied forces (Fig. 3c), loading rates (Fig. 3e), the effective stiffness of beads (the ratio between forces and displacements, Fig. 3f), and recruitment of paxillin to beads (Fig. 3b,g). As previously described^42^, unstimulated beads did not recruit paxillin (Fig. 3b,k). High frequencies led to higher deformation rates (speed), loading rates, bead stiffness, and paxillin recruitment than low frequencies (Fig. 3h-k). However, and unlike in the case of stretch, the response was monotonic, and no decrease in reinforcement or paxillin recruitment was observed even at very high frequencies (Fig. 3k). Applying a square rather than triangular 1 Hz signal dramatically increased force loading rates by almost two orders of magnitude (Fig. 3i). Accordingly, it increased, rather than decreased, paxillin recruitment (Fig. 3k).

**Fig. 3:**
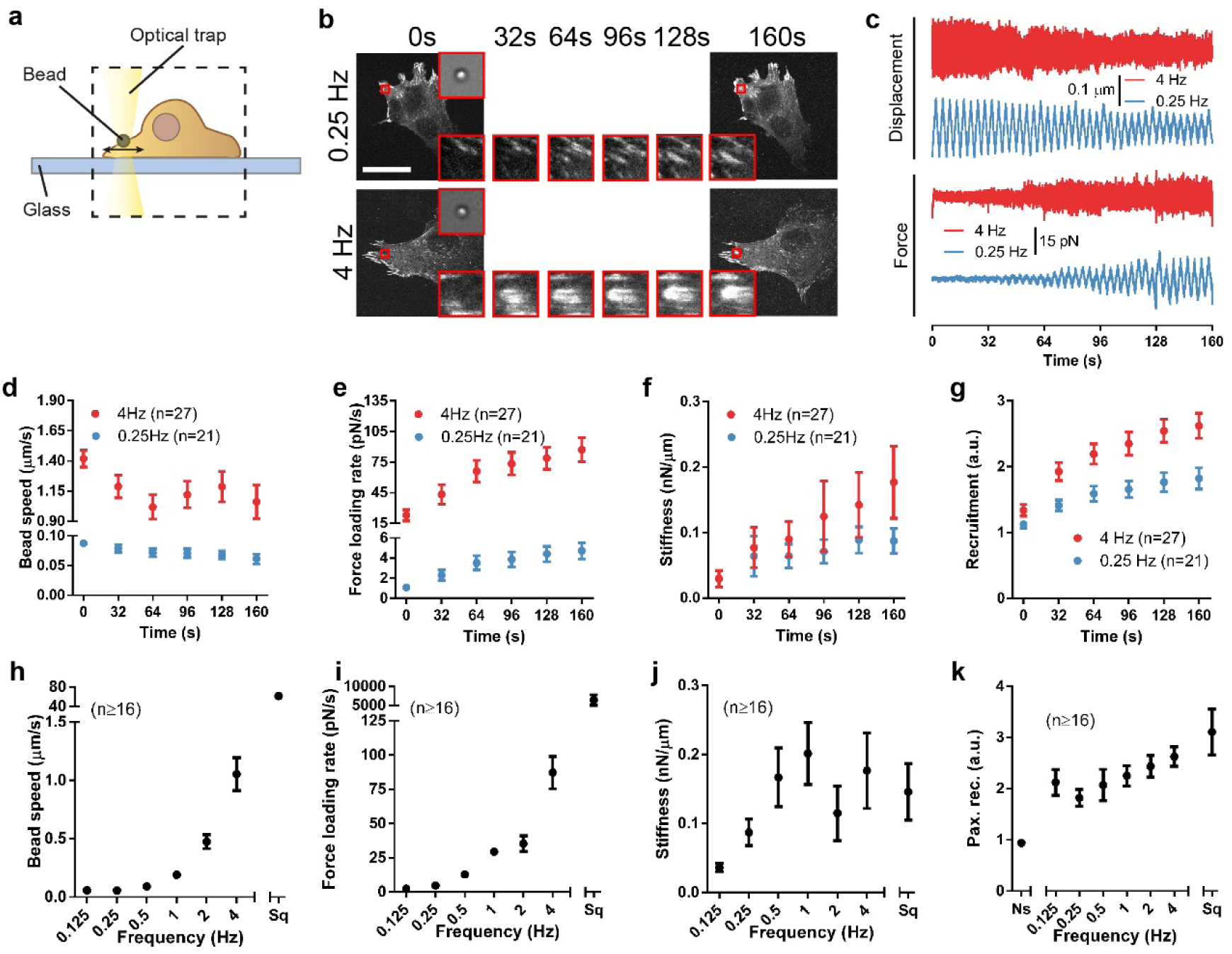
The loading rate of force application to single adhesions drives their maturation. **a**, Illustration of the optical tweezer setup. **b**, Images of cells transfected with GFP-paxillin during force application with triangular signals at 0.25 Hz and 4 Hz, shown as a function of time. The area circled in red indicates the position of the stimulated bead, which is shown magnified at the top-right corner (brightfield image) and bottom-right corner (GFP-paxillin image). Magnified GFP-paxillin images are shown at different timepoints. **c,** Example traces of displacement and forces for beads stimulated at 4 Hz and 0.25 Hz. **d-g**, Bead speed (d), force loading rate (e), stiffness (f), and recruitment of GFP-paxillin at beads (g) as a function of time for beads stimulated at 4 Hz and 0.25 Hz. **h-k,** Bead speed (h), force loading rate (i), stiffness (j), and recruitment of GFP-paxillin at beads (k) for beads at the end of the experiment (160s) for all conditions. The effects of frequency on both stiffness and paxillin recruitment were significant (p=0.002 and 0.008, respectively). Ns, non-stimulated beads, Sq, stimulation with a 1 Hz square signal. n numbers are beads in all panels. Scale bar is 50 µm. Data are shown as mean ± s.e.m.

These results demonstrate that effects happen at the local adhesion scale. Further, they are consistent with a role of the loading rate, which unlike the deformation rate, increased concomitantly with paxillin recruitment both with time (within each experiment) and with frequency. Of note, the absence of the fluidization regime in these experiments is also to be expected, given that optical tweezers cannot apply forces larger than ∼ 100 pN. This is consistent with the observation that cytoskeletal stiffness increased (rather than decreased) with frequency when similarly small forces were applied with magnetic twisting cytometry^43, 44^. Additionally, optical tweezers experiments were carried out in the lamellar region of cells seeded on stiff substrates. This region exhibited stress fibres, and therefore an actin network much more structured than on the rounded cell phenotype found on soft substrates before stretch (Supplementary fig. s3). Thus, forces applied to beads are likely distributed among many filaments, reducing the likelihood of fluidization.

### Fluidization limits mechanosensing at high deformation rates

Our data are consistent with fluidization happening in the case of stretch experiments, but not optical tweezers. To test this experimentally, we attached cells in suspension to a flat Atomic Force Microscope (AFM) cantilever, placed them in contact with a fibronectin-coated glass, and pulled at different speeds (Fig. 4a,b). In these conditions, cells remained rounded, thereby mimicking the low-stiffness phenotype that cells exhibited before being stretched. We then measured the effective stiffness (Young’s modulus) of cells as they were being pulled and thereby stretched (Fig. 4c). Increasing pulling speeds first increased stiffness, as typically occurs in cells or cytoskeletal networks^43–47^. However, between pulling speeds of 5 and 6 μm/s there was a sharp decrease, indicating a partial cytoskeletal disruption, or fluidization. Calculating an approximate equivalence, the range of stretch at which cells failed to mechanosense (10%-20% stretch between 1 and 2 Hz, for cells of ∼ 20 μm in size) corresponds to 2-8 μm/s in deformation, thereby matching the order of magnitude of AFM results (see also Supplementary Table 1). Interestingly, this decrease in stiffness was also associated with lower cell detachment forces, as previously reported^48^ (supplementary fig. 5a). To mimic optical tweezers experiments, we carried out a modified experiment in which we attached fibronectin-coated 3 μm beads to AFM cantilevers, placed them in contact with the surface of previously adhered fibroblasts, and pulled at different speeds (Fig. 4d,e). Consistently with optical tweezers experiments, no softening was observed (Fig. 4f). Of note, measured stiffness values in AFM bead experiments were at least one order of magnitude higher than in whole cell AFM experiments, confirming the notion that cellular lamella have a more structured cytoskeleton than rounded cells (Fig. 4c,f).

**Fig. 4:**
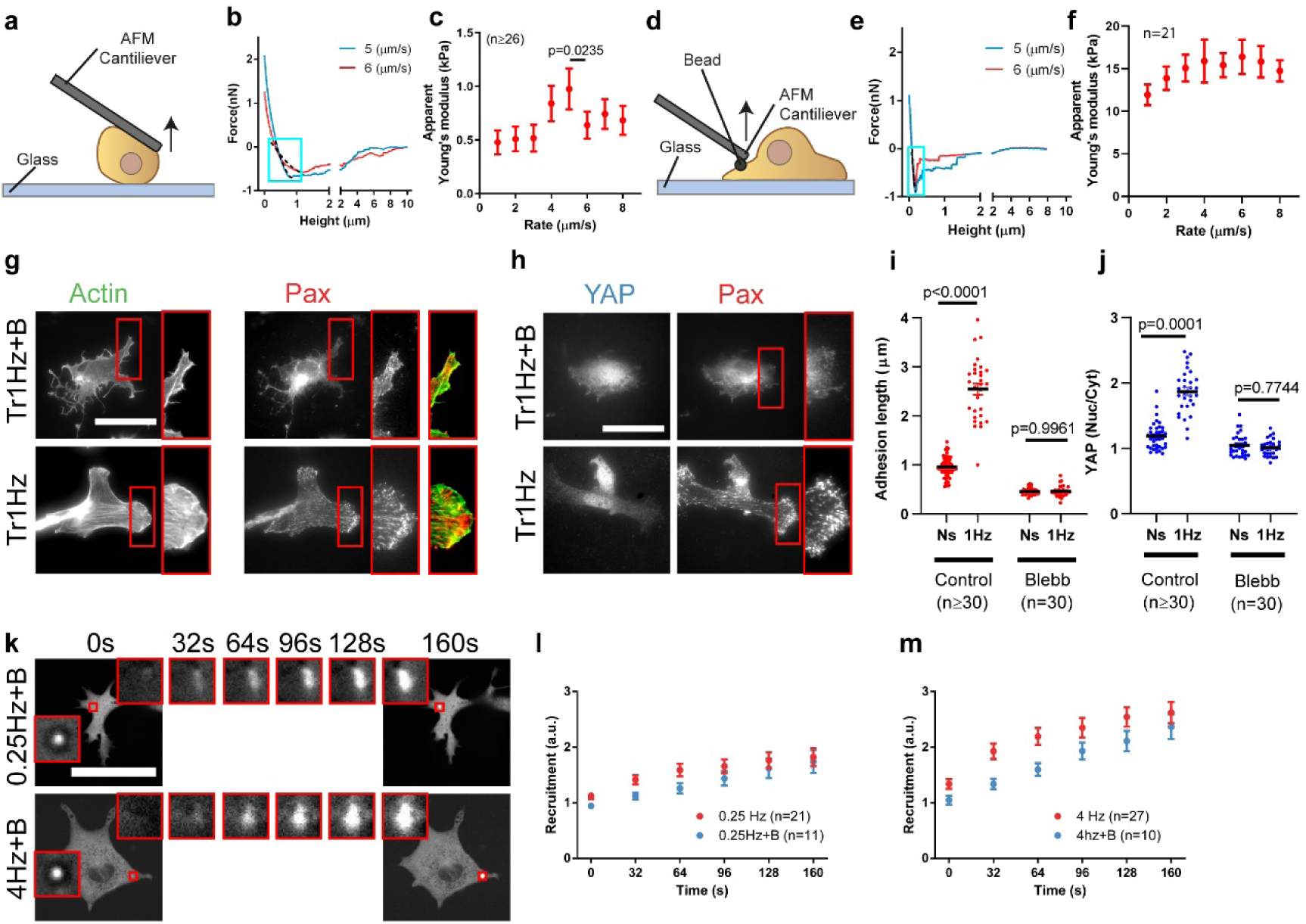
High deformation rates lead to cytoskeletal fluidization. **a**, Illustration of the single cell AFM setup. **b**, Example cantilever retraction curves of the single cell AFM experiments. Dashed lines within blue squares show the fits of the force/deformation curves used to calculate apparent cell stiffness (Young’s modulus). **c**, Stiffness as a function of the retraction speed for cells attaching to a fibronectin-coated substrate. The effect of retraction speed (p<0.0001), and the specific decrease from 5 to 6 µm/s (p=0.0235) were significant. n numbers are curves. **d,** Illustration of the bead AFM setup. **e,** Example cantilever retraction curves of bead AFM experiments. **f,** Stiffness as a function of the retraction speed for fibronectin-coated beads attaching to cells. The effect of retraction speed was not significant. n numbers are curves. **g**,**h** Actin and paxillin (g) and YAP and paxillin (h) stainings for cells stretched by 10% with a triangular 1 Hz signal, with or without blebbistatin treatment. **i,j,** Corresponding quantifications of adhesion length (i) and YAP nuclear to cytosolic ratios (j). n numbers are cells. **k,** Images of blebbistatin-treated cells transfected with GFP-paxillin during force application with triangular signals at 0.25 Hz and 4 Hz, shown as a function of time. Areas circled in red are shown magnified at different timepoints. **l,m,** Corresponding quantifications of recruitment of GFP-paxillin at beads at 0.25 Hz (n) and 4 Hz (o). No significant effect of blebbistatin was observed. n numbers are beads. Scale bar is 50 µm. Data are shown as mean ± s.e.m.

**Fig. 5:**
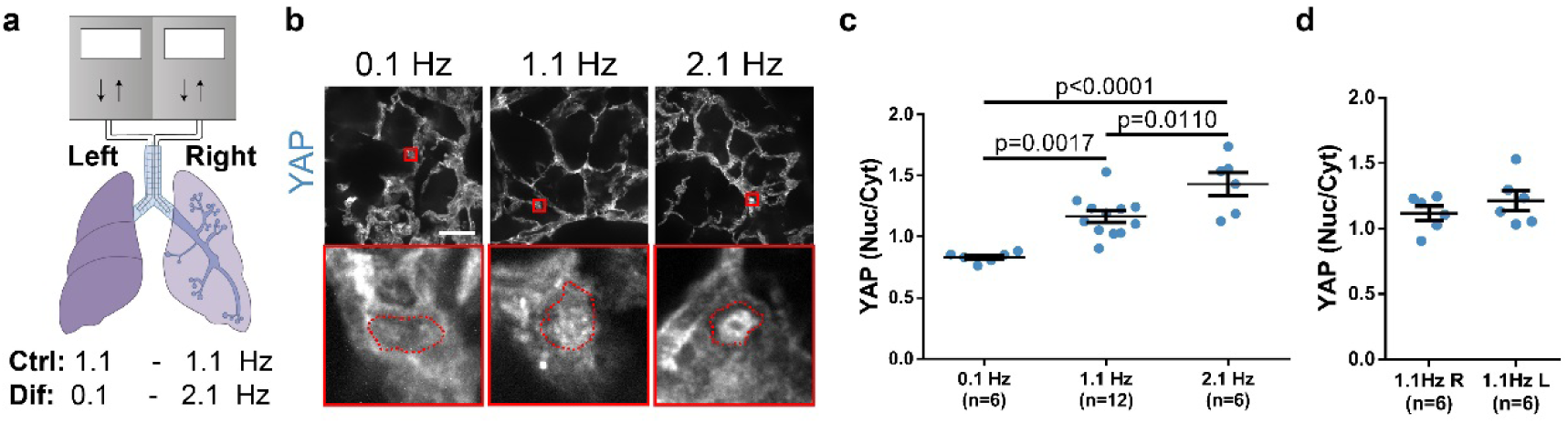
Increasing rates of lung ventilation *in vivo* induce YAP nuclear localization. **a**. Diagram of the rat lung ventilation setup, where each lung was independently cannulated and ventilated. Ventilation was either uniform at 1.1 Hz in both lungs (Ctrl) or differential with the left and right lungs ventilated at 0.1 and 2.1 Hz, respectively (Dif). Both conditions had the same ventilation volume. **b**, YAP staining of rat lungs ventilated at 0.1 Hz, 1.1 Hz, and 2.1 Hz. Areas circled in red are magnified below each image. In magnified images, nuclear contours (as determined from Hoechst stainings) are shown in red. Scale bar is 50 µm. **c**, Quantification of YAP nuclear to cytoplasmic ratios for rat lungs ventilated with same tidal volume at 0.1 Hz, 1.1 Hz, and 2.1 Hz. **d**, Quantification of YAP nuclear to cytoplasmic ratios for left and right rat lungs ventilated at 1.1 Hz. n numbers are rat lungs. No significant differences were observed. Data are shown as mean ± s.e.m.

Finally, we tested another interesting implication of our hypothesis. Loading rates depend not only on the rate of deformation but also on the stiffness of the structure being deformed (which will also impact the amplitude of loads that can be reached). In stretch experiments, the deforming structure is the entire cell, and thus the ability to form stress fibres spanning the cell body (which increases cell stiffness^49^) should be crucial to enable mechanosensing responses. In contrast, in optical tweezers experiments, the presence of a local actin meshwork should be sufficient. To test this, we treated cells with the myosin contractility inhibitor blebbistatin^50^. This affected the actin cytoskeleton by eliminating stress fibres, but still retaining a dendritic actin network in the lamellar region (supplementary fig. 5c,d). In such conditions and as expected, mechanosensing was abrogated in response to stretch (Fig. 4g-j), but not to local force application (Fig. 4k-m).

### Increasing rates of lung ventilation *in vivo* induce YAP nuclear localization

To conclude our study, we assessed whether the role of the force loading rate could also be observed at an organ level *in vivo*. To this end, we used a previously described setup^51^ to subject each of the two lungs of a rat to independent mechanical ventilation (Fig. 5a). This setting allowed us to compare the effects of locally varying the mechanical loading rate in both lungs without interfering with the systemic animal gas exchange. After 1 hour of stimulation, lungs were excised and immunostained, and lung alveoli (containing mostly endothelial and epithelial cells)^52^ were imaged. The 3D, *in vivo* setup led to paxillin immunostainings without sufficient resolution to quantify adhesion shapes, but YAP nuclear to cytosolic ratios could be assessed (Fig. 5b,c, and supplementary Fig. 6). Whereas, as expected, no significant differences were found between right and left lungs when ventilated at the same frequency (Fig. 5d), we found increased levels of nuclear YAP with increased ventilation frequency (Fig. 5b,c and supplementary Fig. 5). These results could not be explained by regulation of oxygen levels, since overall ventilation in the animal was kept constant when differential ventilation was applied to both lungs, and cells in the lung parenchima are perfused with systemic oxygenated blood^53, 54^. Further, hypoxia increases YAP levels^55–57^, as opposed to the effect we see in slowly ventilated (0.1 Hz) lungs. Thus, while live tissues are highly complex environments where neither force transmission nor ensuing signalling cannot be precisely controlled, our results are consistent with a role of loading rate in controlling ventilation-induced mechanotransduction at the organ level in the lungs.

## Discussion

In summary, our results show that force loading rates drive mechanosensing by increasing reinforcement and adhesion growth at the local adhesion level, in a talin-dependent way and as predicted by a molecular clutch model. The range of relevant loading rates at the molecular level is hard to assess, but loading rate dependencies have been predicted to apply over extremely wide ranges^17^, as confirmed for instance in integrin-ECM bonds (10^1^-10^5^ pN/s)^18^. If one assumes a density of bound integrins from 10^0^ to 10^2^ per µm^2^ (from initial to stable adhesions^58, 59^, our range of applied loading rates (Supplementary Table 1) would be largely expected to fall within this range. If loading rates are too high, the cytoskeleton undergoes fluidization, impairing mechanosensing. This fluidization event most likely involves the disruption of actin filaments, since it was prevented by a drug that stabilizes them (Jasplakinolide)^37^. This provides a unifying mechanism to understand how cells respond not only to directly applied forces, but also to passive mechanical stimuli such as tissue rigidity or ECM ligand distribution, where we have reported similar biphasic dependencies of focal adhesions and YAP localization^23^. Further, it also provides a framework to understand how the seemingly opposed concepts of reinforcement and cytoskeletal softening or fluidization, previously analysed within the context of cell rheology^31, 60^, are coupled to drive mechanosensing. Potentially, this framework could be extended to explain mechanosensing mechanisms beyond focal adhesion formation and YAP, such as the actin-dependent nuclear localization of MRTF-A^61^. *In vivo*, the extremely wide range of loading rates in different contexts (from very fast in the respiratory^2^ or cardiovascular^1^ systems, or in vocal cord vibration^62^, to very slow in progressive ECM remodelling in cancer^3^), could thus be central to understand how mechanosensing is regulated. In lung alveoli, we only observed the initial reinforcement phase, even though relevant cell types (fibroblasts, endothelial cells, and epithelial cells) all exhibited both phases in vitro. Whereas the mechanisms remain to be studied, this suggests potential large-scale mechanisms at the tissue level to buffer against cytoskeletal damage. However, in other contexts both reinforcement and fluidization may be at place, and could even be harnessed to establish the levels of mechanical loading required to trigger specific responses.

## Methods

### Cell culture and reagents

Mouse embryonic fibroblasts (MEFs) were cultured as previously described^63^, using Dulbecco’s modified eagle medium (DMEM, Thermofischer Scientific, 41965-039) supplemented with 10% FBS (Thermofischer Scientific, 10270-106) and 1% penicillin-streptomycin (Thermofischer Scientific, 10378-016), and 1.5% HEPES 1M (Sigma Aldrich, H0887). Talin 1-/- MEFs were cultured as previously described^64^, using DMEM supplemented with 15% FBS, 1% penicillin-streptomycin, and 1.5% HEPES 1M. Primary human small airway epithelial cells (SAEC) were purchased from ATCC, cultured in Airway Epithelial Cell Basal Medium (ATCC® PCS-300-030™) supplemented with Bronchial Epithelial Cell Growth Kit (ATCC® PCS-300-040™) and 1% penicillin-streptomycin. Primary human lung microvascular endothelial cells (HMVEC) were purchased from Lonza,cultured using Vascular Cell Basal Medium (ATCC® PCS-100-030), supplemented with Microvascular Endothelial Cell Growth Kit-VEGF (ATCC® PCS-110-041) and 12.5 μg/mL blasticidine. Cell cultures were routinely checked for mycoplasma. CO_2_-independent media was prepared by using CO_2_-independent DMEM (Thermofischer Scientific, 18045 -054) supplemented with 10% FBS, 1% penicillin-streptomycin, 1.5% HEPES 1M, and 2% L-Glutamine (Thermofischer Scientific, 25030-024). Media for optical tweezers experiments was supplemented with Rutin (ThermoFischer Scientific, 132391000) 10 mg/L right before the experiment.

### Transfection

Talin 2 was knocked down as previously described^64^, by transfecting Talin 1-/- MEFs using the Neon Transfection System (Thermofischer Scientific) with a plasmid encoding a short hairpin RNA (shRNA) targeting the nucleotide sequence 5’-GATCCGAAGTCAGTATTACGTTGTTCTCAAGAGAAACAACGTAATACTGACTTCTTTTTTTCTAGAG-3’. For retrograde flow and optical tweezers experiments, cells were transfected with either lifeact-GFP^20^ or pEGFP-Paxillin^65^, using the Nucleofactor 2b Device (Lonza).

### Antibodies and compounds

Primary antibodies used were anti-Paxillin rabbit clonal (Y113, abcam, ab32084), and anti-YAP mouse monoclonal (63.7, Santa Cruz Biotechnology, sc-101199), 1:200. Secondary antibodies used were Alexa Fluor 488 anti-mouse (A-11029, Thermo Fischer Scientific), Alexa Fluor 488 anti-rabbit (A-21206; Thermo Fischer Scientific), and Alexa Fluor 555 anti-rabbit (A-21429, Thermo Fischer Scientific) 1:500. Compounds used were Blebbistatin (Sigma Aldrich) 50 µM, Jasplakinolide (J4580, Sigma Aldrich) 25 nM, phalloidin (Alexa Fluor 555 phalloidin, Thermo Fischer Scientific) 1:1000, and Hoechst (33342, Thermo Fischer Scientific) 1:2000.

For immunofluorescence, the secondary antibodies used were Alexa Fluor 488 anti-rabbit (A-21206; Thermo Fischer Scientific) and Alexa Fluor 555 anti-mouse (A-21422; Thermo Fischer Scientific) diluted 1:200.

### Retrograde flow experiments

Cells were transfected with LifeAct-GFP, plated on gels or glass coated with 10 ug/ml fibronectin, and imaged every 2 seconds for 4 min with a 60× water immersion objective (NA 1.20, Nikon) and a spinning-disc confocal microscope (Andor). For each cell, kymographs were obtained at the cell periphery, and actin speed was measured from the slope of actin features observed in the kymographs.

### Preparation of stretchable membranes

Stretchable polydimethylsiloxane (Sylgard 184 Silicone Elastomer Kit, Dow Corning) membranes were prepared as previously described^66^. A mix of 10:1 base to crosslinker ratio was spun for 1 minute at 500 rpm and cured at 65° C overnight on plastic supports. Once polymerized, membranes were peeled off and assembled onto the stretching device. After assembly, membranes were plasma cleaned for 1 minute, treated with 3-aminopropyl triethoxysilane (APTES, Sigma Aldrich) 10% in ethanol for 1 h at 65 °C, and with glutaraldehyde (Sigma Aldrich) 1.5% in phosphate-buffered saline 1X (PBS, Sigma Aldrich) for 25 min at room temperature.

Then, polyacrylamide gels were prepared and attached to membranes. To this end, polyacrylamide gels were first prepared by adapting previous protocols^23, 67^. Polyacrylamide gels were polymerized between two glass coverslips treated with 2% dimethyldichlorosilane (Plus One Repel Silane, GE Healthcare). For 0.6 kPa gels, the mix contained 4% acrylamide (BioRad), 0.03% BisAcrylamide (BioRad), 2% 200-nm-diameter dark red fluorescence carboxylate-modified beads (Fluospheres, ThermoFischer Scientific), 0.5% ammonium persulphate (APS, Sigma Aldrich), and 0.05% tetramethylethylenediamine (TEMED, Sigma Aldrich), in PBS 1X. After polymerization, one coverslip was detached, and the gel was then attached to the PDMS membrane. To this end, it was pressed against the prepared stretchable PDMS membrane and left overnight at 37 °C in an incubator with humidity control. The remaining coverslip was removed the next day.

Polyacrylamide gels were coated using a protocol adapted from the literature^68^. Briefly, gels were covered with a mix containing 10% HEPES 0.5M Ph 6, 0.002% BisAcrylamide (BioRad), 0.3% 10 mg/ml N-hydroxysuccinimide (NHS, Sigma Aldrich) in dimethyl sulfoxide (DMSO, Sigma Aldrich), 1% Igracure (Sigma Aldrich), 0,0012% tetraacrylate (Sigma Aldrich), in milliQ water. Gels were then covered with a glass coverslip and illuminated with UV light for 10 minutes. After exposure, the glass coverslip was removed, and gels were washed twice with HEPES 25mM Ph 6 and twice again with PBS. Gels were then incubated with 10 µg/ml of fibronectin in PBS overnight at 8°C, washed the next day thrice with PBS and immediately used. The rigidity of the gels was measured using Atomic Force Microscopy as previously described^66^ (see supplementary table 2).

### Cell stretch

Cells were seeded on 0.6 kPa gels attached to previously mounted stretchable PDMS membranes. After attachment, cell media was changed to CO2-independent media. Cells were then stretched at 37°C continuously for 1 hour with one signal type (square or triangular), at one amplitude (20%, 10%, 5%, or 2.5%), and at one frequency (2Hz, 1Hz, 0.5Hz, 0.25Hz, 0.125Hz) to produce different force loading rates. After stimulation, cells were immediately fixed and prepared for immunostaining. To study cell proliferation, cells were stretched using the same protocol for 2.5 hours, and then the Click-iT™ EdU Cell Proliferation Kit (Invitrogen) was used according to manufacturer instructions.

### Immunostainings

Immunostainings were performed as previously described^23^. Cells were fixed with 4% paraformaldehyde for 10 minutes, permeabilized with 0.1% Triton X-100 for 4 minutes, blocked with 2% Fish-Gelatin in PBS 1X for 1 hour, incubated with primary antibody for 1 hour, washed with Fish-Gelatin-PBS for 30 minutes, incubated with secondary antibody for 1 hour, washed with Fish-Gelatin-PBS for 30 minutes, and mounted using ProLong Gold Antifade Mountant (ThermoFischer Scientific).

For immunostainings of animal tissue, at the end-point of experiments animals were sacrificed by exsanguination and during residual heart beating lungs were perfused through the vasculature with ice-cold PBS 1X. Lungs were immediately excised *en bloc* with the heart and intrabronchial cannulas and perfused with cold OCT:PBS (3:1) (Optimal Cutting compound, Company). Lungs were then placed in cassettes with OCT on a dry ice platform and frozen at -80 ° C. Lung blocks 70-µm thick were cut with a cryostat (Thermo Scientific, Massachusetts, MO) and attached to slides. Lung slices were then washed with PBS and fixed with paraformaldehyde 4%. and After three more washes, tissue was permeabilised with 0.2 % Triton X-100, blocked with 10% FBS and incubated for 30 seconds with TrueBlack (Biotium) to reduce ECM autofluorescence. The primary antibody was incubated overnight at 4°C, and after three washes the secondary antibody was incubated during 90 minutes at room temperature. Finally, lung slices were counterstained with NucBlue (Thermo Scientific, Massachusetts, MO) to stain the nuclei and mounted with Fluoromount (Dako). Two lung slices from each lung were imaged in three different fields to a total of six images per condition.

Once the samples were prepared, images of stretched cells were acquired with 60x objective (NIR Apo 60X/WD 2.8, Nikon) with an upright microscope, images of cells on glass were acquired with 60x objective (Plan Apo VC 60X/WD 0.31-0.28, Nikon) with a confocal inverted microscope, and images of animal tissue were acquired with 60x objective (Plan Apo VC Oil 60X/WD 0.13, Nikon) with a confocal inverted microscope.

### Image analysis

Focal adhesion length was quantified manually by assessing the length of three representative adhesions in paxillin stainings at the cell edge, for n cells. Nuclear to cytoplasmic ratio of YAP was quantified manually by segmenting the nucleus using Hoechst (single cells) or NucBlue (rat lung slices) and using the following formula:

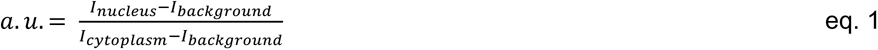

Similarly, protein recruitment to beads was quantified manually by segmenting the bead area using the brightfield image and using the following formula:

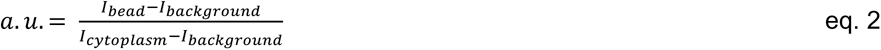

In both cases, I_cytoplasm_ and I_background_ refer to the average fluorescence intensity of the cytoplasm and background (i.e., areas with no cells). I_bead_ and I_nucleus_ refer to the average fluorescence intensity of the nucleus and bead. In the case of rat lung immunostainings, lung cuts were randomized before quantification. Six (6) areas coming from two (2) different cuts were analysed, where twenty (20) cells per area were quantified taking the closest from the geometric centre of the image.

The anisotropy of the actin cytoskeleton was determined by first manually segmenting the cells, followed by analysis using the ImageJ plug-in, FibrilTool^36^.

### Optical Tweezers

The optical tweezers system was adapted from a previous setup^41^. Briefly, the optical tweezer uses a near-infrared fibre laser (λ = 1064 nm, YLR-5-LP; IPG Photonics) that passes through two acoustooptical modulators (DTSX-400-1064; AA Opto-Electronic) which are modulated by a variable frequency driver (Voltage Controlled Oscillator, DRFA10Y2X-D41k-34-50.110; AA Opto-Electronic). After modulation, the beam size is expanded and coupled into an inverted microscope (Eclipse Ti-e; Nikon Corporation) from the rear port. The beam is coupled in the optical path of the microscope by a dichroic mirror and focused in the object plane through a water immersion objective (Plan Apo VC WI, 60x, NA=1.2; Nikon Corporation). To measure the force, the condenser was replaced by a force sensor module (Lunam T-40i; Impetux Optics, S.L.), which was positioned according to the manufacturer procedure. The module is precalibrated and gives direct access to the force applied by the tweezer on any trapped object. To correct for focal drifts during the measurements, the Perfect Focus System (PFS, Nikon Corporation) was used. The sample is heated by a self-built heating chamber, keeping it at 37° C. Image acquisition was done using a spinning disk system (CSU-W1 (Yokogawa); Intelligent Imaging Innovations Inc.) and a CMOS camera (Orca-flash4.0v2; Hamamatsu Photonics K.K.). All hardware was controlled using the custom written LabVIEW programs (National Instruments Corporation).

Beads were coated as described previously^69^. Briefly, carboxylated 1 µm polystyrene beads (Micromod) were coated with a mixture of biotinylated pentameric FN7-10 (a four-domain segment of fibronectin responsible for cell binding and containing the RGD and PHSRN motifs^70^) and biotinylated bovine serum albumin at a ratio of 1:10.

Glass slides were coated with fibronectin 10 ug/ml in PBS overnight and rinsed thrice with PBS. Cells were then seeded on the glass. Once attached, beads were added. One bead was subsequently trapped and placed on the surface of a cell, and the stimulation was then started by using 120 mW of laser power to displace the bead 0.1 µm in x and y from the centre of the trap for 160 seconds by using a triangle or square signal of one frequency (4Hz, 2Hz, 1Hz, 0.5Hz, 0.25Hz, 0.125Hz) to produce different force loading rates. To compensate for the cell dragging the bead towards the nucleus, the optical trap was repositioned at intervals of 32 seconds.

Bead speeds and force loading rates were measured using respectively the displacement and force signals of the optical trap. Using a custom-made MATLAB program, each signal was detrended and then divided into linear segments of individual cycles and fitted to straight lines to obtain the slopes. The stiffness was calculated as described previously^69^ by estimating the transfer function between the force and the displacement data at intervals of 32 seconds, and at the frequency of stimulation.

### Atomic Force Microscopy

AFM experiments were carried out in a Nanowizard 4 AFM (JPK) mounted on top of a Nikon Ti Eclipse microscope. For experiments pulling on beads, Fibronectin or biotin-BSA coated beads were functionalized as described previously^69^. Briefly, carboxylated 3 µm silica beads (Polysciences) were coated with a mixture of biotinylated pentameric FN7-10 (a four-domain segment of fibronectin responsible for cell binding and containing the RGD and PHSRN motifs^70^) and biotinylated bovine serum albumin (Sigma Aldrich) at a ratio of 1:10. The beads where then attached to the cantilevers using a non-fluorescent adhesive (NOA63, Norland Products) to the end of tipless MLCT cantilevers (Veeco). Cells were seeded on fibronectin coated coverslips and a force curve at each retraction velocity was acquired for each of the cells. Cells were kept at 37°C using a BioCell (JPK). The spring constant of the cantilevers was calibrated by thermal tuning using the simple harmonic oscillator model.

For experiments pulling on cells, we followed the protocol described in the literature^48^. Briefly, cantilevers were submerged in sulfuric acid 1M for 1h. They were then washed with milliQ water and plasma cleaned for three minutes. After this, cantilevers were incubated with 0.5 mg/ml biotin-BSA and left overnight in a humid chamber at 37°C. Next day they were washed thrice with PBS and incubated with 0.5 mg/ml Streptavidin and left in a humid chamber at room temperature for 30 minutes. Again, they were washed thrice with PBS and reincubated with 0.4 mg/ml biotin-ConcanavalinA for 30 minutes in a humid chamber at room temperature, after which they were washed thrice with PBS and stored under PBS for use. We used a Nanowizard 4 AFM (JPK) on top of a Nikon Ti Eclipse microscope. ConcanavalinA-coated MLCT-O cantilevers were calibrated by thermal tuning. Then, cells were trypsinized, resuspended in CO_2_ independent media and allowed to recover for 5 minutes. Rounded cells were attached to MLCT cantilevers by exerting 3 nN forces on top of a region of a coverslip with no coating and incubated for 5 minutes. The measurement of the adhesion forces was done by approaching the cell at the same velocity of the corresponding withdraw velocity, keeping for 10 seconds and withdrawing until the cell was fully detached.

The maximum detachment force was determined by analysing the retraction F-D curve. A MATLAB program was used to extract and analyse force-displacement curves from JPK data files. Forces were corrected as described in the literature^71^ by subtracting the baseline offset, i.e. the value obtained at the end of curves when cells were fully detached. We calculated apparent stiffness (Young’s modulus) by fitting retraction curves between the values of zero force and the maximum detachment force F_ad_, defined as the absolute value of the minimum of the curve. This corresponds to the part of the curve from the onset of cell stretching (positive force values at the beginning of curves correspond to cells still in compression) to the point at which cells start detaching. Curves were fitted to the Derjagin, Muller, Toropov (DMT)^72, 73^ contact model, which considers the force-indentation relationship between a flat surface and a sphere, taking into account both elastic deformations and adhesion between the surfaces. Specifically, we fitted the curve:

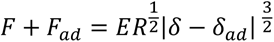

Where F is force, E is Young’s modulus, R is the average radius of cells or beads depending on the experiment (which we took as 10 and 1.5 µm, respectively), δ is indentation, and δ_ad_ is the indentation value corresponding to F_ad_. Note that F_ad_ is defined as positive in this equation.

### Differential rat lung ventilation

Twelve pathogen-free male Sprague Dawley rats (350-450 g) were randomly distributed in the different experimental groups (6 for 2.1Hz-0.1Hz and 6 for 1.1Hz-1.1Hz). Animals were housed in controlled animal quarters under standard light, temperature and humidity exposure. All experimental procedures were approved by the Ethical Committee for Animal Research of the University of Barcelona (Approval number 147/18).

Animals were anesthetized intraperitoneally using 20% urethane (10 mL/kg). After confirmation of deep anaesthesia by tail and paw clamp, animals were tracheostomised and each lung of the rat was independently cannulated (16G; BD Bioscience, San Jose, USA). After muscular relaxation with 0.4 mg/kg pancuronium bromide (Sigma Aldrich, St. Louis, MO) intravenously injected through the penile vein, each lung was connected to a customized small rodent ventilator, with a pressure sensor connected at the entrance of each cannula to monitor animal ventilation according to the conditions explained below. Correct cannulation of each lung was confirmed by opening the chest wall and observing proper independent inflation and deflation of each lung. Animals were ventilated with a tidal volume of 21 ml per kg of animal weight (kg-bw) and with a positive end expiratory pressure of 3 cmH_2_O. Control ventilation was set to a typical frequency of 1.1 Hz with the same tidal volume in each of the two lungs of the rat. To test the effect of varying the ventilation frequency on YAP, a different ventilation frequency was applied to each lung while maintaining the control tidal volume. The left lung was ventilated at 0.1 Hz, and the right lung at 2.1 Hz. In this way, the animal received the same total minute ventilation, hence keeping O_2_ and CO_2_ blood gas levels thereby discarding any systemic effect induced by differential ventilation.

### Computational clutch model

#### Summary of previous clutch model

The computational clutch model was developed by adapting a previously described Monte Carlo simulation, which we implemented to understand cell response to substrate rigidity^20^. Briefly, the model considers an actin filament that can bind to a given number of fibronectin ligands on the matrix *n_f_*, through molecular links (clutches) composed of talin and integrin, with spring constant *k_c_*. Fibronectin molecules are all connected in parallel to the substrate, represented by a spring with constant *k_sub_*. The simulation begins with all clutches disengaged. At each time step, unbound fibronectin molecules can bind integrins according to a loading rate *k_on_* = *k_ont_·d_int_,* where *k_ont_* is the true binding rate characterizing the interaction, and *d_int_* is the density of integrins on the membrane. Fibronectin molecules already bound to an integrin-talin clutch can experience two types of events. First, the integrin-fibronectin bond can unbind according to an unbinding rate *k_off_*, which depends on force as a catch bond as experimentally measured^20, 38^. Second, talin molecules can unfold according to an unfolding rate *k_unf_*, which behaves as a slip bond also as experimentall measured^39^. Because the load on each integrin may be shared between talin and other adaptor molecules, the force used to calculate unfolding is corrected by a factor FR, corresponding to the fraction of integrin-transmitted force experienced by talin.

At the end of each time step, the actin filament moves with respect to the substrate at a speed *v*, and total force on the substrate *F_sub_* is calculated imposing force balance:

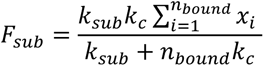

Where *x_i_* is the position of each bound fibronectin molecule, and *n_bound_* is the total number of bound molecules. Further, if a talin molecule has unfolded, mechanosensing is assumed to occur through adhesion growth, which is modelled by increasing integrin density *d_int_* by *d_add_* integrins/µm^2^. If the integrin unbinds before talin unfolding, integrin density decreases by the same amount. However, integrin densities are only allowed to fluctuate between a minimum value *d_min_*, and a maximum value *d_max_*, reflecting the range between nascent adhesions and fully formed focal adhesions.

#### Modifications to model stretch experiments

Beyond this previously implemented model, we modified it in the following ways to specifically reproduce our stretch experiments:

1. First, we introduced a third event (apart from integrin unbinding and talin unfolding) that can occur in a given clutch. This event is the disruption of actin filaments, which had equivalent effects to integrin unbinding (that is, disengagement of the clutch, and reduction in integrin density). This event was only allowed to occur for clutches where talin unfolding (and thereby mechanosensing and reinforcement) had not occurred. We note that although this event is presented as the breaking of actin filaments, it may also include other events disrupting the actin cytoskeleton pulling on clutches, such as for instance disruption of actin crosslinks. Thus, it is not straight forward to assign a given probability rate for the event as a function of force, and for simplicity we simply modelled that this occurred instantaneously above a given threshold of force *F_act_*. The value that best fit our data (142 pN) is in good agreement with experimentally reported values to break actin filaments^40^.
2. Second, the speed *v* of movement between actin and the substrate was not driven by actomyosin contractility (as done in our previous work), but by the applied triangular stretch signals: that is, a constant speed that changed direction twice for each period of the applied signal (with amplitude *A* and frequency *f)*. Thus, |*v*| = 2*Af*, and its sign changed every half cycle. As explained in the main text, the speeds derived from stretch were expected to be similar to actomyosin flows only in the mildest stretch condition (2.5% stretch, 0.125 Hz), so it is reasonable to assume that speeds were mostly driven by stretch across conditions.
3. Third, simulations were not carried out using a fixed time step as done previously, but by a Gillespie algorithm^74^ which only executes one event per time step. That is, for each time step, we calculated stochastically the time at which each possible event for each clutch would occur, according to their respective rates. Then, the algorithm executes only the event occurring first. This was done to ensure that for the highest actin speeds applied (much higher than actomyosin-generated speeds considered in our previous work), potential very fast events were not missed by a fixed, too large time step.
4. Finally and for the same reason, we encountered the problem that for high actin speeds, calculation of rates for the different events were inaccurate. That is, if for instance a given clutch has just been formed and not yet been pulled by actin, its *k_off_* is that corresponding to zero force. However, if actin pulls very quickly, by the time the algorithm predicts an unbinding event, force has risen significantly. This leads to a major change in *k_off_* within the same time step. To account for this, we corrected the different rates to take into account how they were affected by the loading rate. To this end, for each event we first calculated the probability density *p(F,L)* as a function of applied force *F*, when force is ramped starting from zero at a loading rate *L*, as described previously^75^:

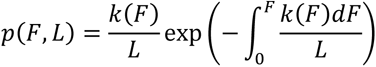

Where *k* is the rate of the event in question (*k_off_* for integrin unbinding or *k_unf_* for talin unfolding). The units of *p* are of force^-1^, and it can be checked that 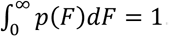. Then, the mean force *F_m_* at which the event takes place can be calculated as:

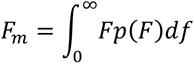

With *F_m_*, one can readily calculate the lifetime of the event as *F_m_*/*L*, and then define a corrected rate k’ as the inverse of the lifetime:

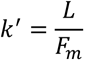

As stated above, this expression applies to the particular case in which force loading starts at zero force, that is *k’* corrects the value of *k*(*F*=*0*) for a given applied loading rate L. In our simulations, at a given time step forces applied to a given clutch are not necessarily zero, and thus we need to consider the case of a force ramping from a given baseline force *F_b_*. To do this, we simply imposed that p(*F*<*F_b_*,*L*) = 0, and normalized the probabilities by 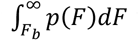 to maintain a total probability of 1.

At the end, instead of the initial rate *k*(*F*) which depends only on force and assumes that force stays constant, we get a corrected rate *k’*(*F*,*L*), which considers that force starts at value *F* and then ramps up with a loading rate *L*. In all cases, we verified that for sufficiently low values of *L*, *k’* = *k*. This confirms the expected result that if the timescale of force loading is slower than that of event lifetime, rates are simply the ones you would obtain by assuming constant force.

Thus, for either *k_off_* or *k_unf_*, calculating this corrected rate allowed to consider the effects of loading rate. For actin breaking events we note that this was not necessary, since we assumed the event to occur instantaneously above a threshold force.

#### Model parameters

All model parameters are described in supplementary table 4. Most parameters were taken exactly as in our previous work, where their choice and relationship to experimental values is discussed^20^. For *k_sub_* (substrate stiffness) we took values corresponding to the lower range employed in our previous work, which match the 0.5 kPa polyacrylamide Young’s modulus used here. Parameters specifically introduced or modified include:

- The frequency (f) and amplitude (A) of the stretch signal applied. Frequencies were chosen to reproduce the experimental range, and amplitude values provided good fits at values ranging from 1.5 to 3 µm. These values are of the order of the expected amplitudes of relative movements of cells ∼ 20 µm in diameter being stretched by 2.5 – 20%. Interestingly, the range of amplitudes providing good fits in the model (1.5-3 µm) is narrower than the experimental range (2.5 – 20%), showing that the experimental sensitivity of cells is less than that predicted by the model.
- A parameter *d_max_* was introduced, setting the maximum value of integrin recruitment (and thereby adhesion growth). This was done to reproduce the experimental range in adhesion sizes. We note that in our previous model, the limit to adhesion growth emerged naturally from the stalling of actomyosin flows, once substrate forces reach the maximum forces that myosin motors can exert. In this model, flow speeds do not stall as they are imposed by stretch, and thus a limit to their growth has to be imposed.
- The number of integrins added each time mechanosensing takes place (*d_add_*) was increased from 10 to 24 /μm^2^ from our previous model. This was necessary to allow the model to respond within the timescale of fast stretch oscillations.
- Finally, as described above a parameter *F_act_* quantifying the force required to break actin filaments was introduced. This value falls within the range of reported experimental values^40^.

Throughout simulations, all parameters except *A* and *f* were fixed, according to the values in supplementary table 4. To model the different experiments, only *A* and *f* were modified, according to the experimental conditions.

#### Model output

Model simulations were run for 1000 s, and then the average values of integrin densities *d_int_* were calculated. These values were then taken as a proxy of adhesion growth, and compared to adhesion length measurements in stretch experiments. To compare the results, model values were scaled between the minimum and maximum experimental length values, and plotted in the same graphs in fig. 2e. Note that this scaling was the same for all panels, and not modified to specifically fit each panel.

### Statistical analysis

No statistical methods were used to determine sample size before execution of the experiments. All independent datasets were first checked for normality using the d’Agostino-Pearson K2 normality test. One-way ANOVA was performed for more than 2 comparisons. Normal one-to-one comparisons were carried out by using a t-test. In case of time-paired data, we used a paired t-test. Multiple comparisons were made using Tukey’s t-test. Non-normal multiple datasets were compared using Kruskal-Wallis’ test, while non-normal two-dataset comparison was done using Man-Whitney’s test. For two factor comparisons the test used was two-way ANOVA, after which Sidak’s multiple comparisons test was performed. All statistic tests were two-sided. All central tendency values are mean and error bars shown are standard error of the mean. Significance is considered for p<0.05. In all cases, at least two independent experimental repeats were carried out for each condition. The exact number of samples (n) is specified in figures or in supplementary table 3.

## Data and code availability

Data are available upon request. Custom scripts are uploaded as supplementary material.

## Acknowledgements

This work was supported by the Spanish Ministry of Science and Innovation (PID2019-110298GB-I00, PGC2018-099645-B-I00), the European Commission (H2020-FETPROACT-01-2016-731957, and the Marie Sklodowska-Curie grant agreement No. 798504 to A.E.A.), the Generalitat de Catalunya (2017-SGR-1602), Fundació la Marató de TV3, the European Research Council (ERC-Adv 883739 to X.T.), the prize “ICREA Academia” for excellence in research to P.R-C., and Obra Social “La Caixa”. S.H and T.B. were supported by the German Science Foundation (EXC 1003 CiM, Cells in Motion), the European Research Council (Consolidator Grants 771201, PolarizeMe) and the Human Frontier Science Program (HFSP grant RGP0018/2017). A.E.M.B. was supported by a Sir Henry Wellcome fellowship (210887/Z/18/Z). We thank V. Gonzàlez-Tarragó for assistance in experiments, set-up implementation, and discussions. We thank the members of P. Roca-Cusachs., X. Trepat, T. Betz, I. Almendros, and R. Farré laboratories for technical assistance and discussions. We thank M. Brandt and D. Navajas for technical assistance and discussions. We thank A. Lahiguera and C. Ureña for discussions.

## Contributions

P.R.-C. conceived the study; I.An., B.F, S.H., A-L.L., R.F., T.B., I.Al., and P.R-C designed the experiments; I.An., B.F., X.Q., and J.K. performed the experiments; I.An., B.F., N.C., X.T., J.K., A.E.M.B, A.E-A., T.B., and P.R-C. analysed the experiments; and P.R-C. performed the modelling work and wrote the manuscript. All authors commented on the manuscript and contributed to it.

## Corresponding authors

Correspondence to Isaac Almendros and Pere Roca-Cusachs.

## Supplementary information

### Supplementary figures and tables

**Supplementary Fig. 1:**
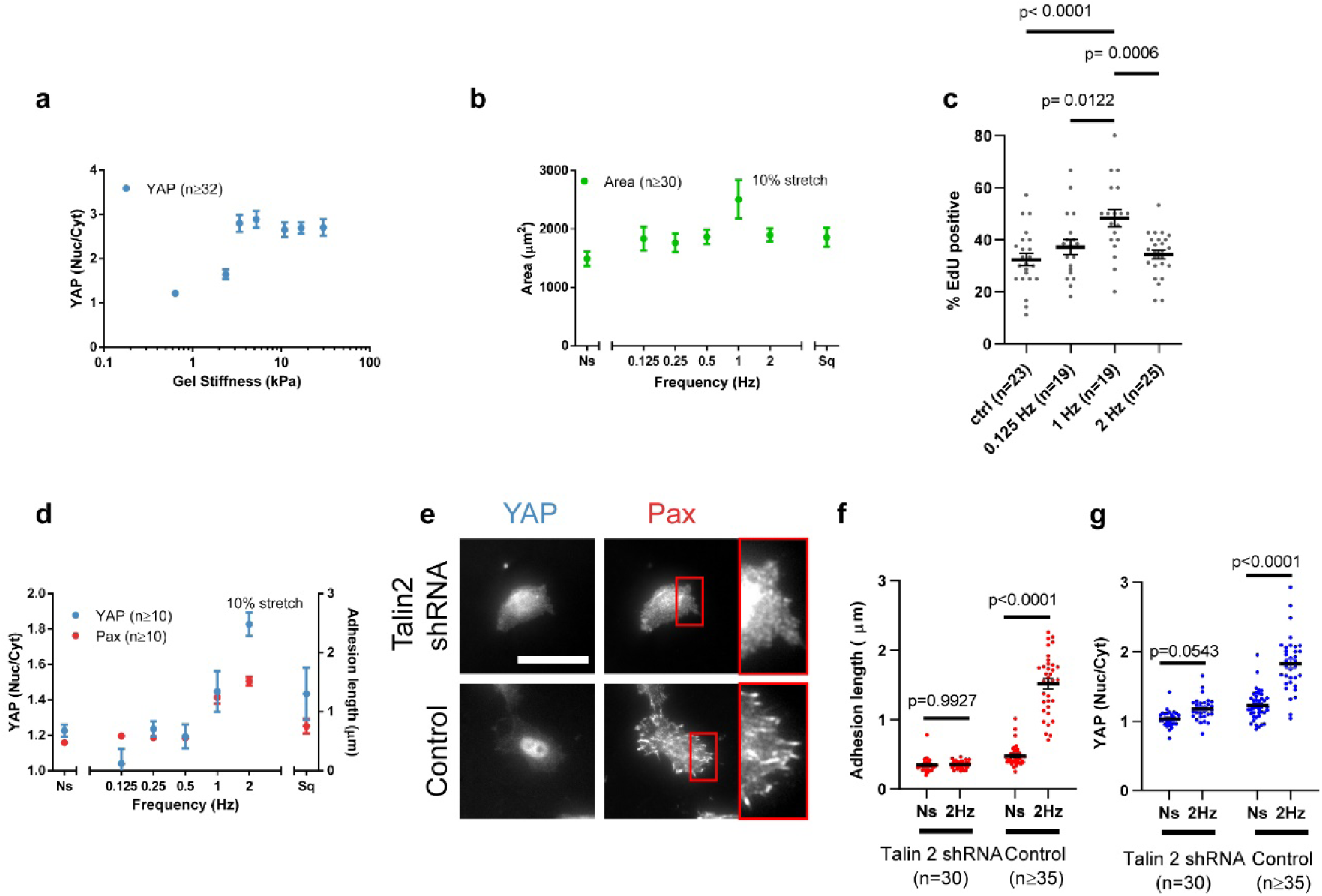
Additional information on stretch experiments. **a,** For mouse embryonic fibroblasts (cells used throughout the paper), quantification of YAP nuclear to cytoplasmic ratios for control cells plated on gels of increasing rigidity. The effect of stiffness was significant (p<0.0001). **b**, Quantification of cell area for cells stretched 10%. The effect of frequency was significant (p=0.0085). **c** For cells stretched by 10%, quantification of cell proliferation rates as a function of stretch frequency, as assessed with an EdU incorporation assay. **d,** Quantification of YAP nuclear to cytoplasmic ratios and focal adhesion lengths for Talin 1 knockdown (control) cells stretched at 10%. These cells overexpress talin 2 and have a wild-type phenotype^20^, and are the control for subsequent talin 2 depletion. The effects of frequency were significant for both YAP and paxillin (p<0.0001). The effect of square versus triangular 1 Hz signals was significant only for paxillin (p=0.0042). **e,** YAP and paxillin stainings of Talin 1 knockdown (control) cells, and Talin 2 shRNA cells stretched at 10%, 2 Hz. In the paxillin image, areas circled in red are shown magnified at the right. These cells showed the peak of response at 2 Hz, and subsequent comparisons were therefore carried out at this frequency. **f,g** Quantifications of focal adhesion lengths (e) and YAP nuclear to cytoplasmic ratios (f) for Talin 1 knockdown (control) cells, and Talin 2 shRNA cells either not stretched (Ns) or stretched at 10% with a frequency of 2 Hz. Scale bar is 30 µm. n numbers are cells in all panels. Data are shown as mean ± s.e.m.

**Supplementary Fig. 2:**
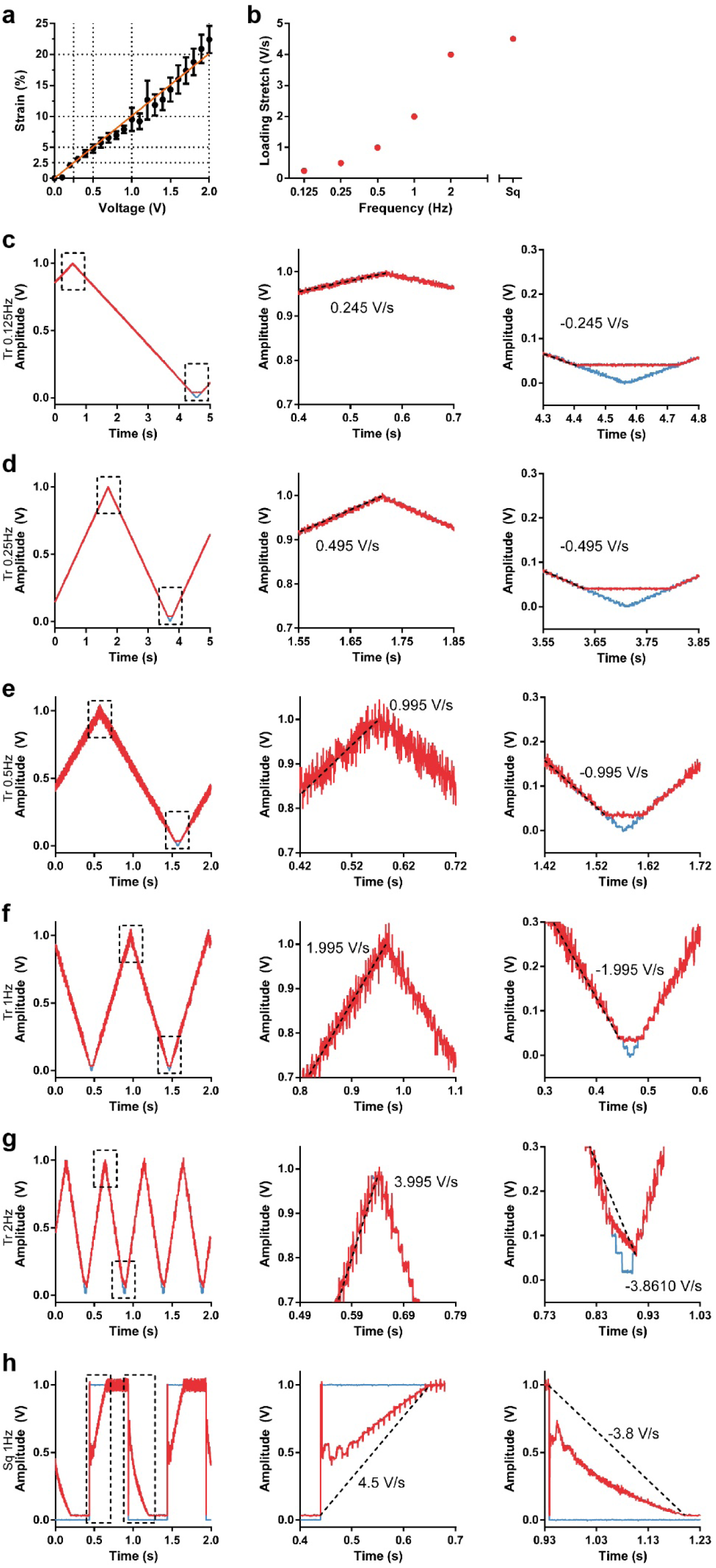
Additional characterization of the cell stretch setup. **a,** Calibration of measured gel stretch as a function of the voltage applied on the pressure transducer. **b**, Applied stretch rate as a function of frequency applied. **c-h,** Example traces of applied and measured voltage signals at the pressure transducer for triangular signals at 0.125 Hz (c), 0.25 Hz (d) 0.5 Hz (e), 1 Hz (f), and 2 Hz (g), and square signals at 1Hz (h). Data are shown as mean ± s.e.m.

**Supplementary Fig. 3:**
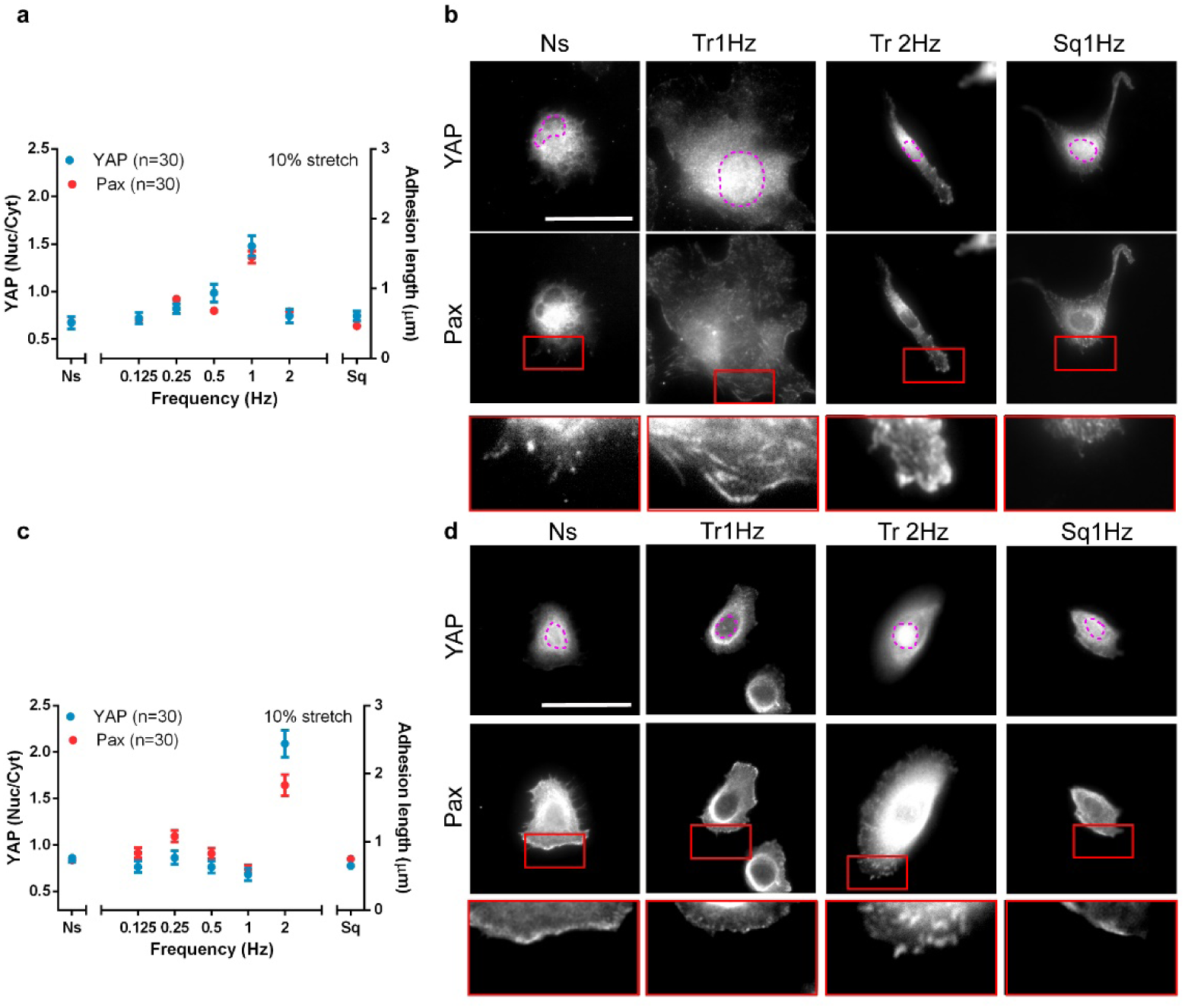
Response to stretch of lung endothelial and epithelial cells. **a**, YAP and paxillin stainings of human microvascular endothelial cells (HMVEC) stretched by 10% using triangular (Tr) and square (Sq) signals at different frequencies. Ns, non-stretched cells. In YAP images, magenta outline indicates the nucleus. In the paxillin image, areas circled in red are shown magnified at the botttom. **b**, Corresponding quantifications of YAP nuclear to cytoplasmic ratios and paxillin focal adhesion lengths. Results are shown for non-stretched cells (Ns), cells stretched with triangular signals at different frequencies, and cells stretched with a square signal at 1 Hz. n numbers are cells. The effects of frequency were significant for both YAP and paxillin (p<0.0001). The effect of square versus triangular 1 Hz signals was significant both YAP and paxillin (p<0.0001). **c**,**d** Same information as in a,b for small airway epithelial cells (SAEC). The effects of frequency were significant for both YAP and paxillin (p<0.0001). The effect of square versus triangular 1 Hz signals was not significant for either YAP or paxillin (p<0.0001). n numbers are cells. Scale bars are 40 μm. Data are shown as mean ± s.e.m.

**Supplementary Fig. 4:**
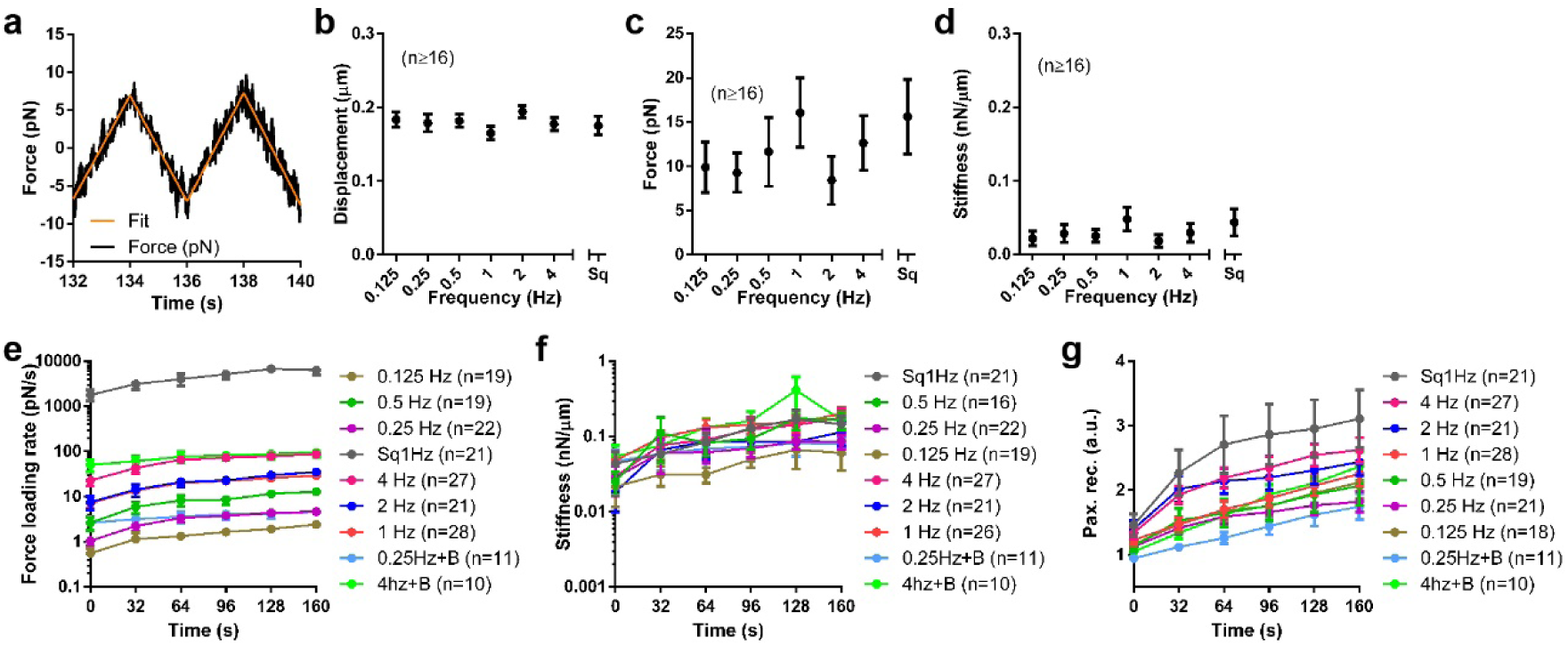
Additional results of the optical tweezers experiments. **a,** Example trace and fit of a measured force signal. To calculate force loading rates, the slope of the fitted lines was taken. **b,** Bead displacement amplitude at time 0s for all frequencies. The effect of frequency was not significant. **c,** Bead force amplitude at time 0s for all frequencies. The effect of frequency was not significant. **d,** Bead stiffness at time 0s for all frequencies. The effect of frequency was not significant. **e,** Force loading rate as a function of time for all conditions. **f,** Stiffness as a function of time for all conditions. **g,** Recruitment of GFP-paxillin to beads as a function of time for all conditions. N numbers are beads in all panels. Data are shown as mean ± s.e.m.

**Supplementary Fig. 5:**
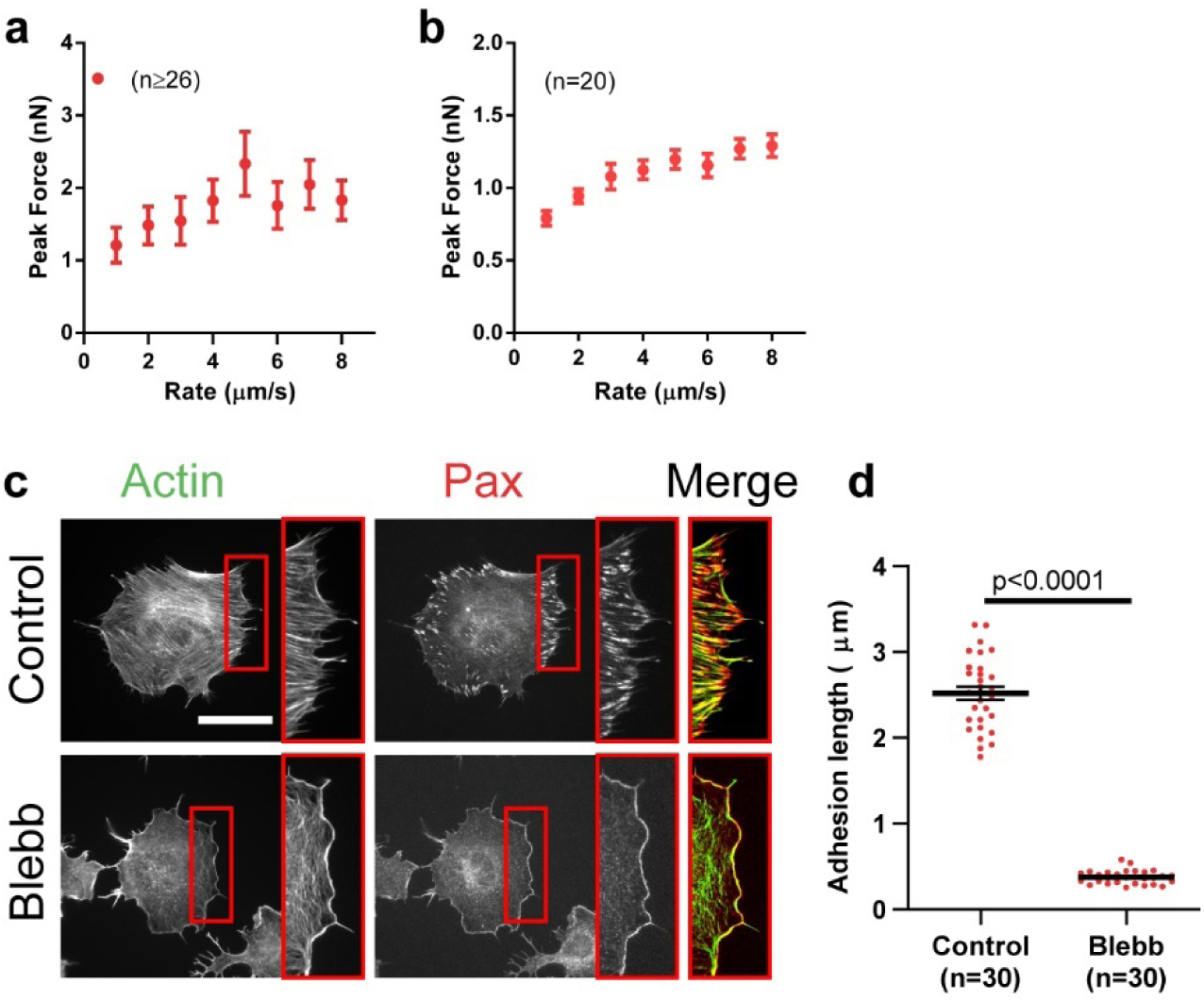
Additional AFM and blebbistatin data. **a**, Peak force during cantilever retraction as a function of the retraction speed for cells attaching to a fibronectin-coated substrate. The effect of retraction speed was significant (p<0.0001). n numbers are curves. **b,** Peak force during cantilever retraction as a function of the retraction speed for fibronectin-coated beads attaching to cells. The effect of retraction speed was significant (p<0.0001). n numbers are curves. **c**, Actin and paxillin stainings in control cells seeded on glass with or without Blebbistatin treatment. Areas circled in red are shown magnified at the right of each image, and shown as a merged image (actin, green, paxillin, red). Blebbistatin disrupts actin stress fibres but not lamellar actin. **d**, Quantification of the adhesion length for control cells, and cells treated with blebbistatin. N numbers are cells. Scale bar is 50 µm. Data are shown as mean ± s.e.m.

**Supplementary Fig. 6:**
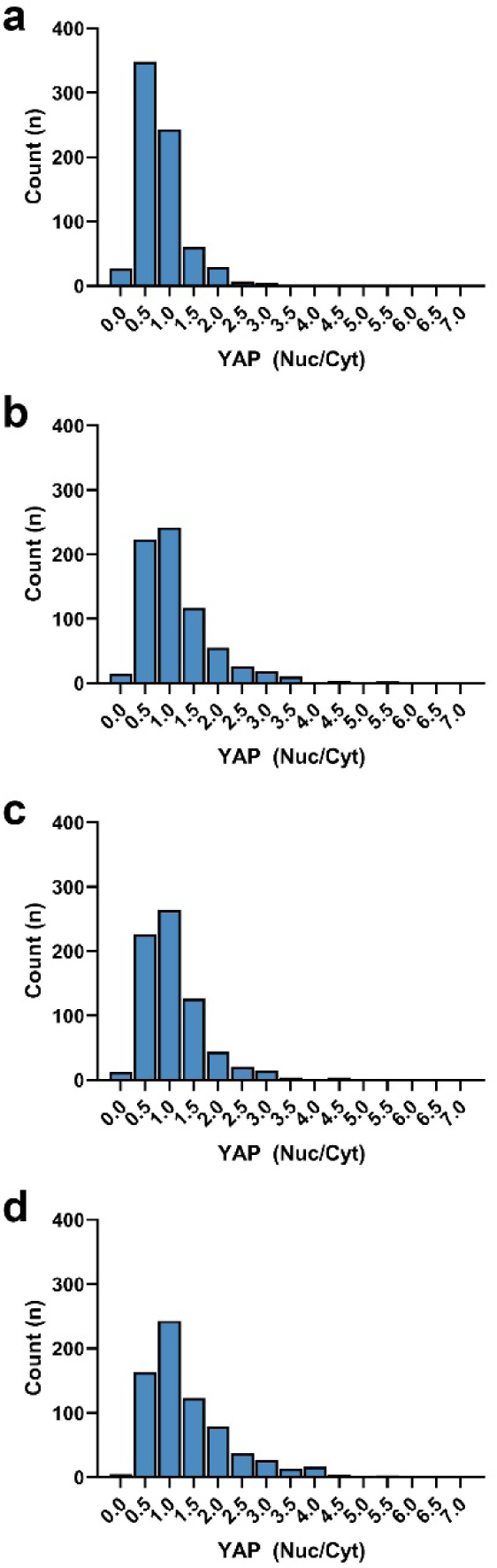
Additional information on rat lung ventilation experiments. Histograms of YAP nuclear to cytoplasmic ratios for rat lung cells ventilated at 0.1 Hz left lung (**a**), 1.1 Hz left lung (**b**), 1.1 Hz right lung (**c**), and 2.1 Hz right lung (**d**). n numbers are cells.

### Supplementary tables

**Supplementary Table 1:** Orders of deformations, loads, and rates in the different figures. Since several of the values are approximate estimates, we provide only orders of magnitude for comparison. (i) Deformations were calculated at the cell edge from stretch applied, assuming an average cell radius of 10 µm. (ii) Loading rates calculated from deformation rates taking stiffness values of the order of 2 nN/µm (fig. 3c). (iii) Cell spreading areas were also estimated from an average 10 µm cell radius, considering that adhesions and forces are largely generated the cell edge (estimated to be of ∼ 2 µm, maximum size of focal adhesions in our measurements). (iv) Average contact areas between 1 µm beads and cells taken from the literature^76, 77^. (v) Contact areas at maximum indentation were estimated to be of ∼5 µm^2^ (bead-cell contact) and ∼25 µm^2^ (cell-substrate contact) by using Hertz contact mechanics^78^.

| Figure | Deformation<br>($\mu\text{m}$) | Def. rate<br>($\mu\text{m/s}$) | Load<br>(nN) | Loading<br>rate<br>(nN/s) | Load per<br>area<br>(pN/$\mu\text{m}^2$) | Loading<br>rate per<br>area<br>(pN/$\mu\text{m}^2/\text{s}$) |
| --- | --- | --- | --- | --- | --- | --- |
| 1<br>(stretch) | $10^{-1} - 10^0$ (i) | $10^{-1} - 10^1$ | $10^0 - 10^1$<br>(ii) | $10^{-1} - 10^1$ | $10^0 - 10^1$ (iii) | $10^0 - 10^2$ |
| 2 (optical<br>tweezers) | $10^{-1}$ | $10^{-2} - 10^1$ | $10^{-2}$ | $10^{-3} - 10^1$ | $10^2$ (iv) | $10^0 - 10^4$ |
| 3c (AFM,<br>cells) | $10^0$ | $10^0 - 10^1$ | $10^0$ | $10^0 - 10^1$ | $10^1$ (v) | $10^1 - 10^3$ |
| 3f (AFM,<br>beads) | $10^{-1}$ | $10^0 - 10^1$ | $10^0$ | $10^1 - 10^2$ | $10^2$ (v) | $10^3 - 10^4$ |

**Supplementary Table 2:**
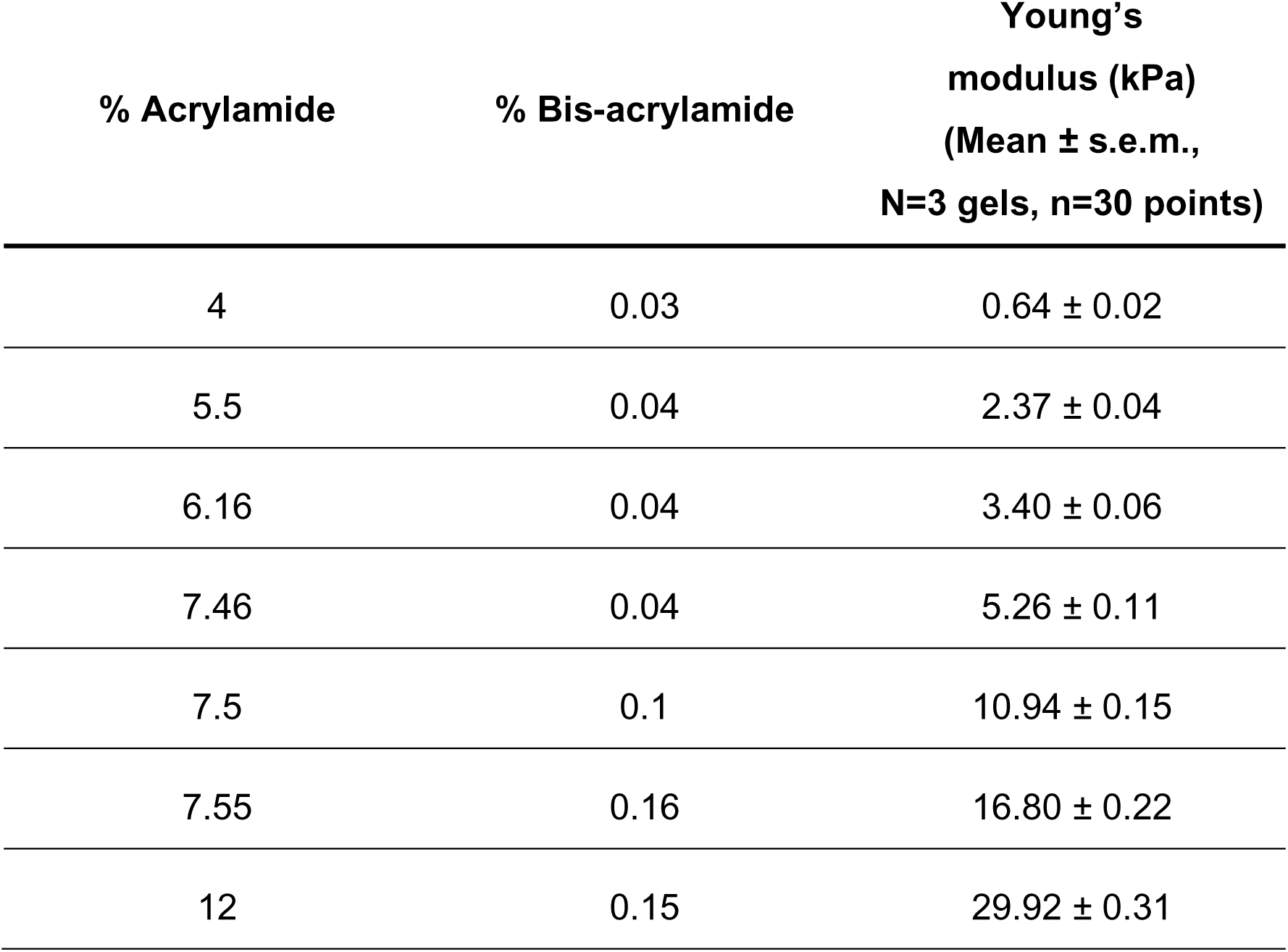
Polyacrylamide gel rigidities measured with AFM.

**Supplementary Table 3:**
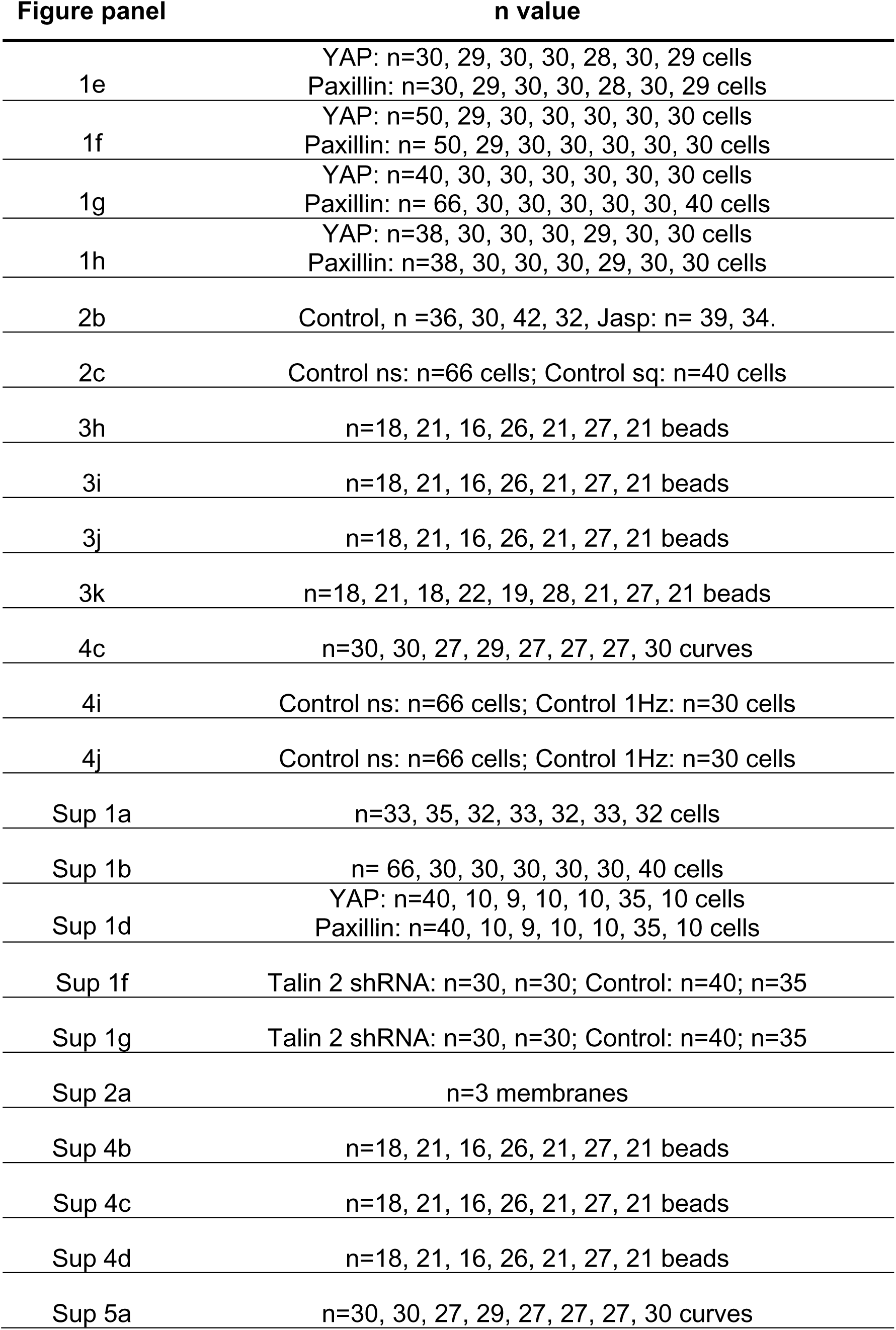
Exact n values for panels where the information did not fit in the figure.

**Supplementary table 4.** Model parameters.

| Parameter | meaning | Value | Origin |
| --- | --- | --- | --- |
| $n_f$ | Number of FN molecules | 1200 | 20 |
| $K_{oni}$ | Intrinsic binding rate | $2.11 \times 10^{-4} \mu\text{m}^2/\text{s}$ | 20 |
| $K_{off}$ | Integrin unbinding rate | Catch bond | 20 |
| $K_{fold}$ | Talin unfolding rate | Slip bond | 20 |
| $k_c$ | Clutch spring constant | 1 nN/nm | 20 |
| FR | Fraction of force experienced by talin | 0.073 | 20 |
| $k_{sub}$ | Substrate spring constant | 1 $\mu\text{N}/\text{nm}$ | Lower range in<sup>20</sup> |
| $d_{min}$ | Minimum (baseline) Integrin density on the membrane | 300 $/\mu\text{m}^2$ | 20 |
| $d_{max}$ | Maximum integrin density on the membrane | 1200 $/\mu\text{m}^2$ | Adjusted |
| $d_{add}$ | Integrins added after each reinforcement event | 24 $/\mu\text{m}^2$ | Adjusted |
| A | Amplitude of stretch signal | 1.5 – 3 $\mu\text{m}$ | Adjusted |
| fr | Frequency of stretch signal | 0.125 Hz – 2 Hz | Experimental values |
| $F_{act}$ | Force for cytoskeletal fluidization | 142 pN | Adjusted |

### Supplementary videos

**Supplementary Video 1:** Time lapse of cell transfected with LifeAct-GFP and plated on a polyacrylamide gel of 0.5 kPa in stiffness.

**Supplementary Video 2:** Time lapse of cell transfected with LifeAct-GFP and plated on fibronectin-coated glass.

